# Beyond the Axenic Culture Filtrate Secretome: BONCAT and conventional proteomics together reveal the Infection-associated secreted protein repertoire of *Acinetobacter baumannii* in the human lung Epithelial cell line A549

**DOI:** 10.64898/2026.10.25.761628

**Authors:** Mohd Ilyas, Ruchi Tanwer, Prabhakar Babele, Niti Singh, Pratibha Sharma, Shweta Tyagi, Yukti Gupta, Sneh Bajpai, Avika Kalra, Mahima Kaushik, Rahul Mahlawat, Saifullah Muzaffar Husain, Pratiksha Pandita, Aastha Dua, Manjistha Baruah, Krishnamohan Atmakuri

**Author notes:** **Correspondence:** Krishnamohan Atmakuri, PhD, Bacterial Pathogenesis Group, Centre for Microbial Research, Translational Health Science and Technology institute, Faridabad, India. **Co-correspondence and First Author:** Mohd Ilyas, PhD, Department of Chemistry and Integrated Science Lab (IceLab), Umeå University, Umeå, SWEDEN. **Author contributions.** Designing research studies; M.I, K.A. Conducting experiments, M.I., R., P.S., S.T., Y.G. Acquiring data; M. I., N.S., R., P.B., P.S., S.T., Y.G. S.B., A. K., M.K., R.M., S., A.D., M.B., P.P. Analysing data; K.A., M. I., N.S., R., P.B., P.S., S.T., Y.G. S.B., A. K., M.K., R.M., S., A.D., M.B., P.P. Writing the manuscript; K.A., M.I., P.B. Editing and finalizing manuscript: K.A., and M.I.

## Abstract

Acinetobacter baumannii (Ab) is an important opportunistic pathogen responsible for pneumonia, bloodstream infections, and sepsis. Although several virulence determinants have been characterized, the infection-associated bacterial protein repertoire associated with host-cell infection remains incompletely understood. Conventional proteomic analyses of axenic cultures provide valuable information but cannot distinguish proteins synthesized before infection from those produced during host–pathogen interactions. This study combined bioorthogonal noncanonical amino acid tagging (BONCAT) with conventional proteomics to characterize the infection-associated Ab’s secreted protein repertoire recovered from human lung epithelial A549 cells.

To achieve pathogen-selective metabolic labelling, an Ab clinical isolate that had caused neonatal sepsis (Ab7) was engineered to express a methionyl-tRNA synthetase variant, MetRSᴺᴸᴸ (L10N, Y258L, H299L), capable of incorporating azidonorleucine (ANL) into newly synthesised bacterial proteins. Labelling conditions were optimized using methionine biosynthesis inhibitory amino acids, which reduced endogenous methionine availability without appreciably affecting bacterial growth. Efficient ANL incorporation was confirmed by flow cytometry, confocal microscopy, biotin–streptavidin assays, and in-gel fluorescence. Metabolic labelling also remained compatible with bacterial intracellular survival.

Sequential labelling with azidohomoalanine (AHA) before infection and ANL during A549 infection enabled the temporal assignment of bacterial protein synthesis. Subsequent SDS–PAGE fractionation and LC–MS/MS identified 152 infection-associated proteins, including 23 containing experimentally detected amino acid analogue modifications. Among these, 13 were AHA-only, nine were ANL-only, and one contained both analogues. Notably, 117 proteins were not detected in either axenic reference proteome. The infection-associated fraction exhibited enrichment of predicted signal peptide-containing and outer-membrane proteins, while functional annotation identified enrichment of bacterial secretion system components among infection-only proteins.

These findings demonstrate the complementary strengths of BONCAT and conventional proteomics: BONCAT provides evidence of bacterial protein synthesis during infection, whereas conventional proteomics expands detection beyond proteins containing identifiable analogue-modified peptides. Although extracellular recovery does not establish active secretion, the combined approach reveals a host-associated bacterial protein repertoire extending beyond axenic observations. This framework supports future investigations into bacterial adaptation, host–pathogen interactions, and the prioritization of candidate diagnostic biomarkers, subject to functional characterization and clinical validation. The study establishes a methodological foundation for investigating infection-associated protein synthesis and release in complex host–pathogen environments

## Introduction

To thrive within the host, bacterial pathogens deploy a multifaceted virulence arsenal that enables them to sense and respond to hostile environments, evade immune defences, replicate efficiently, and establish protected niches for survival (1–5). Portions of this arsenal are (i) actively secreted and (ii) passively sloughed off into the host milieu, where they reprogram cellular pathways and host cell functions, dampen hosts’ protective responses, and promote pathogens’ persistence and their dissemination (6–8). *Acinetobacter baumannii* (Ab), a leading opportunistic Gram-negative pathogen responsible for blood stream infections, sepsis, pneumonia, meningitis, and urinary tract infections, also exploits the above strategies to colonize, survive and establish a niche within their hosts (2,9–14)

Given Ab’s propensity to trigger pneumonia and sepsis, especially among those in Intensive Care Units, the host pulmonary epithelium is a critical interface for Ab adhesion, invasion, persistence, modulation, and disease establishment (10,15,16). Importantly, disruption of the alveolar epithelial barrier during severe pulmonary infection may facilitate bacterial dissemination into the bloodstream, contributing to bacteremia and potentially sepsis, thereby underscoring the relevance of identifying Ab’s proteins associated with the lung epithelial cellular milieu. Hence, we exploited the human alveolar epithelial cell line A549 as the *in vitro* platform, particularly to investigate Ab’s infection-associated secreted protein repertoire that may facilitate epithelial colonisation, intracellular survival, and host-cell injury. It is well established that Ab exists both as an intercellular and intracellular pathogen at different stages of infection and its spread in the host (17,18).

Extensive research has uncovered several facets of Ab’s biology, persistence, and virulence mechanisms (3,7,8,18). For instance, components of its cytosolic and surface-associated protein repertoire such as OmpA, YjjK, VgrG, and EF-Tu have been implicated in cell adhesion, host invasion, and effectors’ delivery (3,19–21). Additionally, proteins including LipA, LipH, CpaA, and InvL are known to subvert diverse host processes and contribute to Ab-induced pathology (6,22,23). Beyond these surface-anchored or cytosolic factors, Ab utilizes specialized secretion systems to export virulence determinants directly into its host environment. These include: (i) Type I secretion system-mediated delivery of Bap to promote adherence and biofilm maturation; (ii) Type II secretion system export of LipAN for efficient lung colonization; (iii) Type V secretion system secretion of Ata for influencing virulence; (iv) Type VI system-mediated release of ppApp synthetase and RhS factors for driving interbacterial antagonism; and (v) Outer membrane-derived extracellular vesicles-mediated delivery of OmpA and CsuA for reinforcing virulence and biofilm development (1,19,21,22,22,24).

Although these important virulence determinants have been identified through classical molecular microbiology approaches, molecular genetics, targeted mutagenesis, comparative genomics, and *in vitro* secretion studies (22,25,26), our understanding of infection associated Ab protein during *ex vivo* or *in vivo* infection remains limited. Mass spectrometric-based proteomic analyses of axenic bacterial cultures and their extracellular milieu provide an essential baseline but cannot identify the repertoire of bacterial proteins released into or associated with host epithelial cells during infection. Therefore, identifying and characterizing the infection associated Ab protein is critical for advancing our understanding of Ab-mediated pathogenesis and identifying potential therapeutic targets (26). Importantly, emerging evidence suggests that Ab possesses a dynamic, condition-responsive infection associated Ab protein that extends beyond established virulence determinants and may undergo substantial remodelling in response to the host cellular environment.

Uncovering the infection-associated secreted proteome of Ab presents considerable technical challenges. Traditional reporter fusions and N- or C-terminal protein-tagging strategies can reveal bacterial proteins that reach the A549 cellular milieu; however, the introduction of such tags may alter protein localization, secretion efficiency, or biological function, thereby limiting their broader applicability (27). Conventional liquid chromatography–tandem mass spectrometry (LC–MS/MS)-based proteomics provides an unbiased approach to identifying proteins within complex biological samples but cannot distinguish newly synthesized proteins from those synthesized before infection (28,29). Moreover, the overwhelming abundance and diversity of host-derived proteins can obscure low-abundance bacterial proteins, limiting their identification. Although protein fractionation by SDS–PAGE followed by in-gel digestion and LC–MS/MS can improve proteome coverage and facilitate the detection of otherwise underrepresented bacterial proteins, this approach does not provide information on nascently synthesised proteins. Thus, conventional proteomics is best complemented by approaches that selectively label and enrich newly synthesised bacterial proteins during infection, enabling a more comprehensive characterisation of the Ab protein repertoire associated with the host cellular milieu.

To overcome the limitations of conventional proteomics, we employed bioorthogonal non-canonical amino acid tagging (BONCAT) to selectively label newly synthesized Ab proteins during infection of human lung epithelial cells (A549) (29,30). BONCAT relies on the incorporation of non-canonical amino acids (ncAAs) bearing bioorthogonal functional groups, such as azides or alkynes, into newly synthesized proteins. These functional groups subsequently undergo selective click-chemistry reactions with complementary probes, enabling the enrichment, visualisation, and identification of labelled proteins, even within complex biological samples dominated by host-derived proteins (refs). When combined with pathogen-selective metabolic labelling, BONCAT facilitates the preferential enrichment and identification of newly synthesised bacterial proteins during infection, including those that may escape detection by conventional proteomic approaches.

Under physiological conditions, methionyl-tRNA synthetase (MetRS; also referred to as methionyl-tRNA ligase, MetG) catalyses the attachment of L-methionine (L-Met) to its cognate tRNA. Although MetRS exhibits high substrate specificity, certain methionine analogues, including L-homopropargylglycine (HPG) and L-azidohomoalanine (AHA), can be incorporated into newly synthesized proteins by endogenous MetRS enzymes in both bacterial and mammalian cells (29,31–34). To achieve pathogen-selective labelling during infection, we engineered an Ab strain expressing a MetRS variant, MetRSᴺᴸᴸ (L10N, Y258L, H299L), capable of incorporating azidonorleucine (ANL), a bioorthogonal methionine analogue that is not efficiently utilised by wild-type MetRS. By integrating BONCAT-mediated metabolic labeling with conventional SDS–PAGE-based LC–MS/MS proteomics, we sought to comprehensively characterize the infection-associated Ab protein repertoire within the A549 cellular milieu. This complementary strategy extends detection of bacterial proteins beyond those recovered from axenic cultures and provides a foundation for identifying candidate virulence-associated proteins and for understanding Ab–host interactions.

## Materials and Methods

### Primary sequence alignment of *E. coli* and Ab Methionyl-tRNA synthetases and superimposition of their active site secondary structures

Protein sequences of *E. coli* Methionyl-tRNA synthetase (MetRS; also referred to as Methionyl-tRNA ligase or as MetG; PDB ID: 1PG2) and Ab MetRS (MetG, PDB ID: 5URB) were aligned using ClustalW (reference). The corresponding ribbon structures were superimposed using WinCoot (reference). The L-Methionine binding site amino acid residues (Leu10, Tyr258 and His299 in Ab MetRS) were found to be conserved in *E. coli* MetRS as Leu13, Tyr260, and His301. The deposited 5URB sequence is identical to Ab7 MetRS across all 546 residues it models.

### Bacterial strains and their growth conditions

Clinical isolates of Ab designated as #7 (Ab7) and #755 (Ab755) (from our laboratory collection), along with the reference strain AB5075 (University of Washington, USA), were routinely cultured in Luria–Bertani (LB; Miller formulation; HiMedia, India) broth or agar at 37 °C. When required, bacteria were sub-cultured in M9 minimal medium (35). The composition of 1× M9 broth was as follows: 0.1 mM CaCl₂ (Merck–Sigma Aldrich), 2 mM MgSO₄ (Merck–Sigma Aldrich), 47.74 mM Na₂HPO₄·7H₂O (HiMedia), 22.04 mM KH₂PO₄ (SRL, India), 8.55 mM NaCl (Merck–Sigma Aldrich), and 18.69 mM NH₄Cl (HiMedia), supplemented with 0.4% succinate (Merck–Sigma Aldrich) as the sole carbon source.

*E. coli* strain NEB 5-alpha (New England Biolabs, USA) was cultivated in LB broth or agar (HiMedia, India) at 37 °C (35). The following antibiotic concentrations were employed: for Ab, Gentamicin 25 µg/mL (HiMedia) and Colistin 8 µg/mL (Thermo Fisher Scientific, USA); for *E. coli*, Chloramphenicol 25 µg/mL (Sigma Aldrich), Hygromycin 100 µg/mL (Invitrogen, USA)Kanamycin 100 µg/mL (Sigma Aldrich), and Carbenicillin 100 µg/mL (Merck–Sigma Aldrich).

### Cloning and site-directed mutagenesis

The Ab wild-type *metRS* (*metRS*^LYH^) gene was used as the template to generate the *metRS*ᴺᴸᴸ (L10N, Y258L, H299L) variant. The generated variant construct was cloned into a compatible shuttle plasmid under the control of an arabinose-inducible promoter (L-(+)-arabinose). The *metRS*ᴺᴸᴸ variant was generated in three successive steps: (i) L10N substitution by PCR-based cloning, (ii) Y258L substitution via site-directed mutagenesis, and (iii) H299L substitution by PCR-based cloning. To generate *metRS*ᴸ¹⁰ᴺ, PCR was performed using Phusion High-Fidelity polymerase (New England Biolabs, USA), genomic DNA (gDNA) from Ab7 as template, and primers KAP600 (forward, harboring L10N) and KAP543 (reverse) (Table S1). The amplified variant amplicon (∼2064 bp) was Gateway-cloned into pDONR221 vector (Thermo Fisher Scientific, USA). Two rounds of Gateway cloning were conducted using primer sets KAP542/KAP543 (first round) and KAP266/KAP267 (second round) (Table S1). The recombinant *metRS*ᴸ¹⁰ᴺ amplicon was recombined into pDONR221 using BP Clonase II (as per manufacturer’s protocol), yielding construct pMI1 (Table S2). The L10N substitution was confirmed via Sanger sequencing (Applied Biosystems 3500 Genetic Analyzer; Thermo Fisher Scientific).

To introduce Y258L, site-directed mutagenesis was performed on pMI1 using primers KAP601 (forward) and KAP602 (reverse) with Phusion High-Fidelity polymerase (NEB). Following mutagenic PCR, the parental template (pMI1) was digested with DpnI for 90 min, and the mutated plasmid DNA (*metRS*ᴺᴸ) purified using a PCR purification kit (MDI, India). The purified plasmid was transformed into chemically competent *E. coli* NEB 5-alpha, yielding construct pMI2 (pDONR221 + *metRS*ᴺᴸ; Table S2). Successful insertion of Y258L was also confirmed by Sanger sequencing.

The H299L substitution was introduced into *metRS*ᴺᴸ in two steps. First, the N-terminal 1–907 bp fragment of *metRS*ᴺᴸ (pMI2 template) was amplified using primers KAP600 (forward) and KAP654 (reverse), the latter harboring the H299L substitution and an internal HaeIII restriction site. After sequence confirmation, the HaeIII-digested 907 bp fragment (bearing L10N, Y258L, and H299L) was stored. In parallel, the full-length *metRSᴺᴸ* from pMI2 was amplified using KAP600 (forward) and KAP591 (reverse, with an XbaI site at the 3′ end), followed by digestion with HaeIII and XbaI. The resulting 1157 bp (908–2064 bp) fragment was ligated with the stored 907 bp fragment to reconstruct the full-length *metRS*ᴺᴸᴸ, which was cloned into EcoRV/XbaI-digested pBlueScript KS+, yielding pMI3. The complete variant gene was then amplified with primers KAP590 (forward, with XhoI overhang) and KAP591 (reverse, with XbaI overhang), digested with XhoI/XbaI, and cloned into pVRL2 (a shuttle plasmid replicative in Ab;(36), to produce pMI4 harboring *metRS*ᴺᴸᴸ under an arabinose-inducible promoter.

### *E. coli* and Ab transformation

Transformation of the Ab7 was performed following a modified electroporation protocol (37). Briefly, 100 mL of Ab7 culture was grown at 37 °C and 200 rpm to an OD₆₀₀ of 0.8, chilled on ice for 30 min, and centrifuged at 3200 × g, 4 °C. Cells were then washed thrice with 100 mL sterile, ice-cold 10 % glycerol and finally resuspended in 6 mL of 10 % glycerol. Aliquots (100 µL) were stored at −80 °C. For electroporation, aliquots were thawed on ice, mixed gently with 250–300 ng plasmid DNA, and transferred into pre-chilled 2 mm cuvettes (Bio-Rad, USA). Electroporation was performed using a Gene Pulser (Bio-Rad) set to 1.8 kV, 200 Ω, and 25 µF. Cells were immediately recovered in 1 mL sterile SOC medium (HiMedia) and incubated for 1 h at 37 °C, 220 rpm. Transformants were selected by plating 100 µL of the recovered culture on LB agar containing the appropriate antibiotics and incubating overnight at 37 °C. All *E. coli* transformations were performed using chemically competent cells (38).

### Ab growth in rich, minimal, and non-canonical amino acid (ncAA)-containing media

Ab7 cultures were routinely revived from −80 °C glycerol stocks on LB agar with required antibiotics. Primary cultures were grown in 5 mL LB broth at 200 rpm, 37 °C. Mid-log-phase cells were pelleted (3200 × g, 25 °C, 10 min), washed thrice with sterile LB (rich) or M9 (minimal; 0.4 % succinate as carbon source) broth, and sub-cultured at OD₆₀₀ = 0.005 in the same medium. When cultures reached OD₆₀₀ ≈ 0.2–0.3, cells were washed again (3200 × g, 25 °C, 10 min) and re-inoculated (OD₆₀₀ = 0.005) for growth monitoring. Growth kinetics were assessed by plotting OD₆₀₀ and CFU/mL against time.

When required, Azidohomoalanine (AHA; Click Chemistry Tools, USA) and Azidonorleucine (ANL; Amatek Scientific, China) were added to M9 medium to final concentrations of 0.25 mM and 1 mM, respectively. Stock solutions (1 M) were prepared in sterile double-distilled water and filtered through 0.22 µm filters (MDI, India).

Where indicated, M9 medium was supplemented with sodium acetate (8 mM; reference) or MBIAA (L-Methionine-biosynthesis-inhibitor amino acids), L-Lysine, L-Threonine, and L-Phenylalanine each at 100 µg/mL final concentration, and L-Isoleucine, L-Leucine, and L-Valine each at 50 µg/mL, added at OD₆₀₀ = 0.1 for 16 h (39). Stocks of sodium acetate (5 M) and individual amino acids (50 mg/mL for L-Lysine, L-Valine, L-Threonine; 10 mg/mL for L-Leucine, L-Phenylalanine, L-Isoleucine) were prepared in sterile double-distilled water and filter-sterilised (0.22 µm; MDI, India).

### Metabolic labelling of proteomes in Ab and its recombinant derivatives

Wild-type Ab7 and its recombinant derivative expressing the engineered methionyl-tRNA synthetase MetRSᴺᴸᴸ (Ab7 pMI4) were cultured in M9 medium supplemented with 0.4% succinate, as described above. Cultures were inoculated at an initial OD₆₀₀ of 0.005 and grown to OD₆₀₀ approximately 0.1. At this optical density, arabinose was added to Ab7 pMI4 cultures at a final concentration of 2% to induce MetRSᴺᴸᴸ expression. Arabinose (HiMedia, India) was prepared as a 20% stock solution in sterile water and filter-sterilised through a 0.22 µm membrane. For metabolic labelling, AHA or ANL was added to the designated cultures at final concentrations of 0.25 mM or 1 mM, respectively. Cultures were incubated with the respective analogue for 12 h before harvesting or preparation for infection, as appropriate to the experiment. Where indicated, cultures were also exposed to MBIAA, the amino-acid mixture described above, for 12 h.

### Evaluation of ncAA incorporation into proteomes of Ab and its engineered NLL variant

#### Surface labelling

Approximately 10⁸ AHA- or ANL-labelled bacterial cells were washed twice (3200 × g, 4 °C, 10 min) with sterile 1× PBS (pH 7.4). Pellets were resuspended in 200 µL PBS (pH 7.4) and treated for 30 min in the dark with 40 mM iodoacetamide to block free cysteine –SH groups and prevent nonspecific DIBO-alkyne binding. Cells were then washed three times with 500 µL PBS and resuspended in 100 µL PBS (pH 7.4) containing 15 µM DIBO-alkyne-conjugated Alexa Fluor 488 (stock 1 mM in DMSO; sub-stock 15 µM in PBS). Labelling was carried out for 1 h at RT in dark conditions with shaking (300 rpm; Thermomixer Comfort, Eppendorf, USA). Cells were washed thrice (3200 × g, 4 °C, 10 min) with 1 mL PBS to remove unbound fluorophore, filtered through 5 µm syringe filters (MDI), and gently passed three times through a 26 G needle to obtain a single-cell suspension. Efficiency of ncAA incorporation was assessed by FACS, confocal microscopy, and in-gel fluorescence.

#### FACS analysis

A 20 µL aliquot (∼2 × 10⁵ cells) of Alexa Fluor 488-labelled bacteria was diluted in 5 mL PBS (pH 7.4) and analysed on a BD FACS Canto II (BD Biosciences, USA). A minimum of 10⁵ events was recorded, and percent fluorescence-positive cells quantified using FlowJo software.

### Confocal microscopy

The remaining 980 µL sample was concentrated to 50 µL by centrifugation, and 10 µL was fixed (10 min, 4 % paraformaldehyde), washed thrice with PBS and mounted in 10 µL mounting medium (20 mM Tris-HCl pH 8.0, 0.5 % N-propyl gallate, 50 % glycerol). Fluorescence imaging was performed using an Olympus FV3000 confocal microscope (Tokyo, Japan) at 60× magnification, and images processed with FV10-ASW v2.01.03.10 software.

### In-gel fluorescence

Click-reagent stock solutions were prepared as follows: 100 mM THPTA ligand (Vector Labs, USA), 20 mM CuSO₄ (Sigma Aldrich), 300 mM sodium ascorbate (Sigma Aldrich), all in sterile water, and 1.5 mM TAMRA-alkyne in DMSO. Sodium ascorbate and CuSO₄ stocks were prepared freshly. Click reactions were carried out in 1.5 mL microcentrifuge tubes containing 80 µg of azide-modified protein lysates. To each tube, 10 µM TAMRA-alkyne, 5 mM THPTA, and 1 mM CuSO₄ were added in sequence and volume adjusted to 100 µL with PBS. After vortexing, 15 mM sodium ascorbate was added to initiate the reaction, which was incubated for 1 h at RT in the dark. Proteins were precipitated by methanol–chloroform method (40) to remove excess fluorophore, air-dried, and resuspended in Laemmli sample buffer with 5 % β-mercaptoethanol. Samples were boiled (95 °C, 15 min) and resolved on 15 % SDS-PAGE (Bio-Rad, USA). TAMRA-labelled proteins were imaged on an Azure Biosystems c400 gel scanner (Cy3 channel 526 nm excitation / 565 nm emission); protein marker bands were visualised using Cy5 channel (∼650/670 nm).

### Semi-quantitation of incorporated ncAAs on the surface proteome of Ab

Semi-quantitative assessment of ncAA incorporation was carried out using biotin-DIBO alkyne labelling. After blocking thiols with 40 mM iodoacetamide (200 µL reaction volume in PBS, pH 7.4, 30 min), cells (1 × 10⁸) were washed thrice with PBS and incubated in 100 µL PBS containing 20 µM biotin-DIBO alkyne (stock 50 mM in DMSO) for 1 h at RT in the dark with shaking (300 rpm). Cells were washed three times with 1 mL PBS to remove unbound biotin-DIBO.

Bound biotin was detected by incubating cells for 2 h on a rocker with streptavidin–HRP (1:4000; 2.5 mg/mL) in 1 mL PBS (pH 7.4). Following three washes (3200 × g, RT, 10 min each), equal numbers of cells from three biological replicates were used for TMB substrate assay (BD Biosciences; (41). Briefly, cells were incubated with 200 µL TMB substrate (1:1 TMB/H₂O₂) for 2 min, centrifuged (3200 × g, RT, 5 min), and 100 µL supernatant transferred to a 96-well plate. The reaction was terminated with 5 % H₂SO₄, and absorbance recorded at 450 nm (Microplate Reader, Bio-Rad) for comparative quantification.

### L-Methionine quantitation

Primary cultures of Ab 7were grown to mid-log phase in LB broth, washed twice (3200 × g, 4 °C, 10 min) in M9 broth, and passaged twice at OD₆₀₀ = 0.005 in M9. When second-passage cultures reached OD₆₀₀ ≈ 0.1, cells were exposed to sodium acetate (8 mM) or MBIAA for varying durations (8, 12, and 16 h) as described earlier. Equal numbers of cells were harvested at each time point, washed twice with PBS (3200 × g, 4 °C, 10 min), and processed for L-methionine quantitation.One millilitre of chilled 80 % methanol was added to each pellet and vortexed for 10 min at room temperature, followed by sonication (40 kHz, 10 min; Branson, USA) and incubation on ice (10 min). Lysates were centrifuged (15,000 × g, 4 °C, 20 min), and 900 µL of supernatant was transferred to fresh microcentrifuge tubes and speed-vacuum-dried (SPD2010-220, Thermo Fisher). The dried residue was reconstituted in 100 µL 50 % acetonitrile, vortexed (5 min, RT), and centrifuged again (15,000 × g, 4 °C, 20 min). Approximately 80 µL of the supernatant was subjected to targeted metabolomics. Bacterial samples were analysed in four biological replicates. A pooled quality-control (QC) sample was prepared by combining 10 µL aliquots from each bacterial biological replicate and included in the analysis. Data were acquired on an Orbitrap Fusion mass spectrometer (Thermo Scientific) and raw data were aligned, peak-picked, and annotated in Progenesis Qi (Non-Linear Dynamics), then normalised, log-transformed, and scaled using MetaboAnalyst 5.0 for statistical analyses.

### *In vitro* infection of A549 cells with *Ab* clinical isolates and derivatives

Human alveolar epithelial A549 cells were cultured as adherent monolayers in DMEM (HiMedia) supplemented with 10 % heat-inactivated fetal bovine serum (HiMedia), 1× MEM essential amino acids (Gibco), 1× non-essential amino acids (HiMedia), 1X sodium pyruvate (Gibco), and 1X GlutaMAX (Gibco). When cells reached 70–80% confluence, they were detached by 10-min incubation in cold PBS, scraped, and counted using a haemocytometer (Olympus CKX41, Japan). Cells were seeded at 2 × 10⁵ cells/well in 24-well plates and 2 × 10⁶ cells/flask in T-75 flasks for infection studies. Bacterial inocula were prepared from cultures as described previously, washed in sterile DMEM, and adjusted to OD₆₀₀ for an MOI of 1:1000. After 3 h of infection at 37 °C, 5 % CO₂, extracellular bacteria were eliminated using colistin (8 µg/mL) for 1 h. At specified time points, epithelial cells were lysed by 30-min incubation in cold sterile water (4 °C; reference). Serial dilutions of lysates were plated on LB agar for CFU enumeration of intracellular bacteria.

### Detection of ANL-incorporated Ab in A549 cells via BONCAT

For in situ fluorescent detection, A549 monolayers (80 % confluent) grown on poly-L-lysine (0.1 %)-coated coverslips in DMEM containing 1 mM ANL were infected with Ab7 expressing *metRSᴺᴸᴸ*. After 3 h, extracellular bacteria were removed by colistin treatment (8 µg/mL, 1 h). Cells were washed thrice with PBS, fixed and permeabilised using the BD Cytofix/Cytoperm Kit (BD Biosciences) as per manufacturer’s protocol.

Strain-promoted alkyne–azide cycloaddition (SPAAC) was performed by incubating cells with 15 µM Alexa Fluor 488 DIBO-alkyne in 2 mL PBS for 1 h at RTin the dark. Coverslips were washed thrice with PBS and mounted using medium containing DAPI (Molecular Probes, Thermo Fisher). Imaging was performed under 60× objective using an Olympus FV1000 confocal microscope (Tokyo, Japan) and processed with FV10-ASW v2.01.03.10 software.

### Sample preparation for tandem mass spectrometry (MS/MS)

#### Axenic bacterial pellet and culture-filtrate proteins

Mid-log Ab7 cultures grown in M9 medium (OD₆₀₀ ≈ 1.0) were centrifuged (3200 × g, 4 °C, 10 min). Pellets were washed thrice with PBS and resuspended in 400 µL bead-beating buffer (PBS + 1× protease inhibitor cocktail; Thermo Fisher). Cells were lysed using a Biospec bead-beater (4 cycles, 30 s each, 1-min cooling on ice between cycles). Protein concentrations were determined by BCA assay, and aliquots mixed with 4× Laemmli buffer (Green & Sambrook, 2012), 5 % β-mercaptoethanol, and boiled (95 °C, 15 min).

Spent culture media were filtered (0.22 µm, Merck Millipore) and proteins precipitated using ammonium sulphate (100 %, 228.3 g per 300 mL). Precipitates were collected by centrifugation (18,000 × g, 4 °C, 20 min), dissolved in 10 mL PBS, and dialysed overnight at 4 °C against 1 L PBS using SnakeSkin™ tubing (3.5 kDa cut-off; Thermo Scientific). Dialysed proteins were concentrated to 200 µL using 3 kDa Centricon concentrators (Millipore) and quantified by BCA assay (Pierce).

#### Post-infection pathogen infection associated Ab protein proteins

Mid-log Ab7 expressing *metRSᴺᴸᴸ* were harvested, washed thrice with M9, and sub-cultured to OD₆₀₀ = 0.005. At OD₆₀₀ ≈ 0.1, cells were incubated with 0.25 mM AHA and MBIAA for 12 h, and metRSᴺᴸᴸ induced with 2 % arabinose over the same 12 h period. Cells were washed thrice with PBS, resuspended in 5 mL L-methionine-deprived DMEM, and ∼10⁸ labelled bacteria were used to infect 10⁵ A549 cells (MOI 1:1000) in L-methionine-free DMEM containing 1 mM ANL.

After 3 h of infection, the A549 monolayers were washed thrice with PBS and extracellular bacteria were killed with colistin (8 µg/mL, 1 h). The monolayers were then washed thrice with PBS and incubated further (9 h, 37 °C, 5 % CO₂) in fresh L-methionine-free DMEM containing 1 mM ANL, giving a total ANL labelling period of 12 h; ANL was absent during the 1 h colistin step. Infected A549 cells were lysed in 200 µL chilled water containing 1× protease inhibitor cocktail (Thermo Fisher), incubated (30 min, 4 °C), and centrifuged (3200 × g, 4 °C, 10 min). The supernatant was filtered (0.2 µm, MDI) to remove bacteria and quantified using the BCA method (Pierce).

#### In-solution trypsin digestion

For the unlabelled axenic bacterial cell-lysate and culture-filtrate samples, 100 µg protein was denatured in 50 µL 8 M urea (4 °C, 30 min), reduced with 5 µL 200 mM dithiothreitol (DTT) at 37 °C for 1 h, and alkylated with 20 µL 200 mM iodoacetamide (IAA) for 1 h at RTin the dark. The mixture was diluted below 2 M urea with 50 mM ammonium bicarbonate (ABC) and digested with trypsin (1:50; Promega, USA) overnight at 37 °C. Peptide mixtures were acidified with 1 % formic acid and desalted prior to LC-MS.

#### In-gel trypsin digestion

Unlabelled axenic bacterial cell-lysate and culture-filtrate samples were analysed by both in-solution and in-gel digestion, whereas the labelled infection-associated fraction recovered from infected A549 cells was analysed by in-gel digestion only.

In-gel digestions were performed as previously described (reference) with minor modifications. Briefly, 30 µg of protein from each sample was resolved on 12 % SDS-PAGE, stained with Coomassie Brilliant Blue R-250, and each lane excised into 12 gel pieces. Bands were destained (50 % acetonitrile, 50 mM ABC; 200 µL; 2000 rpm, 30 min, RT), reduced (10 mM DTT), alkylated (20 mM IAA), and digested overnight with 100 ng trypsin (Promega) at 37 °C. Peptides were extracted using 50 % acetonitrile with 1 % formic acid.

#### LC–MS/MS analysis

Peptide separation was performed using an Eksigent microLC system coupled to a TripleTOF 5600 mass spectrometer (Sciex, USA). Samples were loaded on a trap column (C18-CL, 300 µm × 0.5 mm, 3 µm, 120 Å) and separated on a C18 analytical column (300 µm × 15 mm, 3 µm, 120 Å) with mobile phases A: water/acetonitrile/formic acid (98/2/0.2 %) and B: acetonitrile/water/formic acid (98/2/0.2 %). Flow rate was 5 µL/min. Gradient: 3 % B (2 min), 3–25 % B (40 min), 25–35 % B (10 min), 80 % B (5 min hold), then re-equilibration to 3 % B (3 min).

Data-dependent acquisition was performed with the following parameters: ion source 2300 V, curtain gas 25 psi, nebuliser gas 20 psi, heater gas 10 psi, temperature 150 °C, MS resolution ≥30,000 (FWHM), survey scan 250 ms, and 30 MS/MS scans (100–1500 m/z, threshold 150 cps, charge state 2+–5+). Total cycle time was 2.35 s. Collision energy (CE) was auto-controlled using the IDA CE script with 5 eV spread. Dynamic exclusion was set to 3 s.

#### Database search for protein identification

Protein identification was performed using ProteinPilot v5.0 and the Paragon algorithm (42), against the Ab UniProt database (July 2023, 3799 entries), with Parameters: sample type, identification; cysteine alkylation, iodoacetamide; digestion, trypsin; instrument, TripleTOF 5600; special factors, BONCAT workflow; ID focus, biological modifications; search effort, thorough ID. False discovery rates were estimated against a reversed decoy database, and peptide and protein identifications accepted below 1% FDR. As of September 2026, 2,409 of the 2,990 accessions (80.6%), including all 23 labelled proteins, have been removed from UniProtKB and retained in UniParc (Supplementary Table S3).

#### Data processing and exclusion criteria

Peptide-spectrum matches carrying a mass shift inconsistent with the experimental design were excluded; these arise from the broad modification space searched under the biological-modifications focus (Supplementary Table S6). Because the database excluded the host proteome, eleven identifications with peptide sequences identical to human (n = 10) or Escherichia coli (n = 1) proteins had been assigned to entries annotated as *A. baumannii;* these were treated as host-derived or contaminant and excluded (Supplementary Table S6); none carried an analogue. Peptide rows lacking a confidence value, peptides whose accession was absent from the protein-level report, and proteins left without peptide evidence were also removed. Peptide counts are unique sequences at 90% confidence or above.

#### Protein grouping

To avoid counting one protein more than once through redundant database entries, accessions sharing an identical peptide of seven or more residues were merged into protein groups, with isoleucine and leucine treated as equivalent; shorter sequences were not used, as they occur by chance in unrelated proteins. A group counted as detected in a condition if any member accession had peptide evidence there, and axenic detections combine in-gel and in-solution analyses. All protein counts are groups: the 152 infection associated Ab protein accessions in Supplementary Data 1 correspond to 151 groups, two accessions of one acyl carrier protein having been merged. Peptide counts are unaffected.

Classification and specificity controls Proteins were classified pre-infection if they carried only Met->Aha peptides, infection-phase if only Met->Anl, and continuous if both; tagged (+tag) variants counted as the corresponding analogue. Background rates of false assignment were estimated from the unlabelled axenic samples, filtered by the criteria above so that control and test data were treated identically. Only the two in-gel datasets were used, contributing 213 (axenic bacterial culture filtrate) and 615 (axenic bacterial cell lysate) methionine-containing peptides. The in-solution datasets were excluded from this control, since their searches returned no AHA, ANL or excluded-modification assignments among approximately 32,000 peptides, indicating that these modifications were absent from the in-solution search space; an absence of assignment there is therefore uninformative. Both were retained for protein detection (Supplementary Table S6). Proportions were compared by Fisher exact tests and 95% confidence intervals calculated by the Clopper-Pearson method.

#### Functional annotation

Sequences for accessions removed from UniProtKB were retrieved from UniParc, giving one sequence for each of the 2,990 accessions. Functional annotation was assigned from sequence with eggNOG-mapper v3.0.0-beta6 and eggNOG database v7.0.0 in DIAMOND mode (43–45), which assigns each query to an orthologous group and transfers the KEGG orthology and pathway annotations. Pathway annotation was obtained for 957 of 2,052 detected groups and 46 of the 117 infection-only groups.

#### Enrichment testing

Enrichment was assessed at protein-group level against all annotated detected groups as background, a group counting as carrying a term if any member accession carried it. Terms present in fewer than three groups of the test set were not tested, since single-protein terms yield uninterpretable fold values; this threshold reduced 79 pathways observed in the infection-only set to 19 tested. Pathways were tested by one-sided Fisher exact test, with p values corrected across all tested pathways by the Benjamini-Hochberg procedure and q < 0.05 considered significant. Gene Ontology terms were annotated but not analysed: because eggNOG-mapper transfers annotation from orthologous group members, eukaryote-specific terms were applied to these bacterial proteins, making enrichment uninterpretable. Sequence-derived annotations, namely Pfam domains, KEGG orthology, signal peptides and predicted localisation, are not subject to this transfer.

#### Signal peptides and localisation

Signal peptides were predicted with SignalP 6.0 from the N-terminal 70 residues (46), and subcellular localisation with PSORTb v3.0.2 in Gram-negative mode from full-length sequences (47). A group counted as signal-peptide positive if any member accession received a non-OTHER prediction and was assigned the most specific non-cytoplasmic localisation among its members. PSORTb assigned only prokaryotic compartments across all 2,990 sequences. Proportions were compared by two-sided Fisher exact tests.

## RESULTS

### In silico analysis shows Ab MetRS L-Met binding pocket structurally conserved with *E. coli* MetRS

Aminoacyl-tRNA synthetases catalyze the attachment of specific amino acids to their cognate tRNAs. Although methionine analogues such as L-homopropargylglycine (HPG) and L-azidohomoalanine (AHA) can be incorporated into newly synthesized proteins by endogenous methionyl-tRNA synthetases (MetRS), their incorporation into both bacterial and mammalian proteins by native MetRS limits their utility for cell-selective metabolic labelling (Fig. 1A). To overcome this limitation, Tanrikulu et al. (2009) (48) engineered an *Escherichia coli* (*E. coli*) MetRS variant, MetRSᴺᴸᴸ, through mutagenesis of three residues within the L-methionine (L-Met)-binding pocket (L13N, Y260L, and H301L). This variant enabled the incorporation of L-azidonorleucine (ANL) into newly synthesized proteins, providing a strategy for selectively labelling bacterial proteins in mixed bacterial populations or during host–pathogen interactions (Fig. 1A).

**Figure 1.**
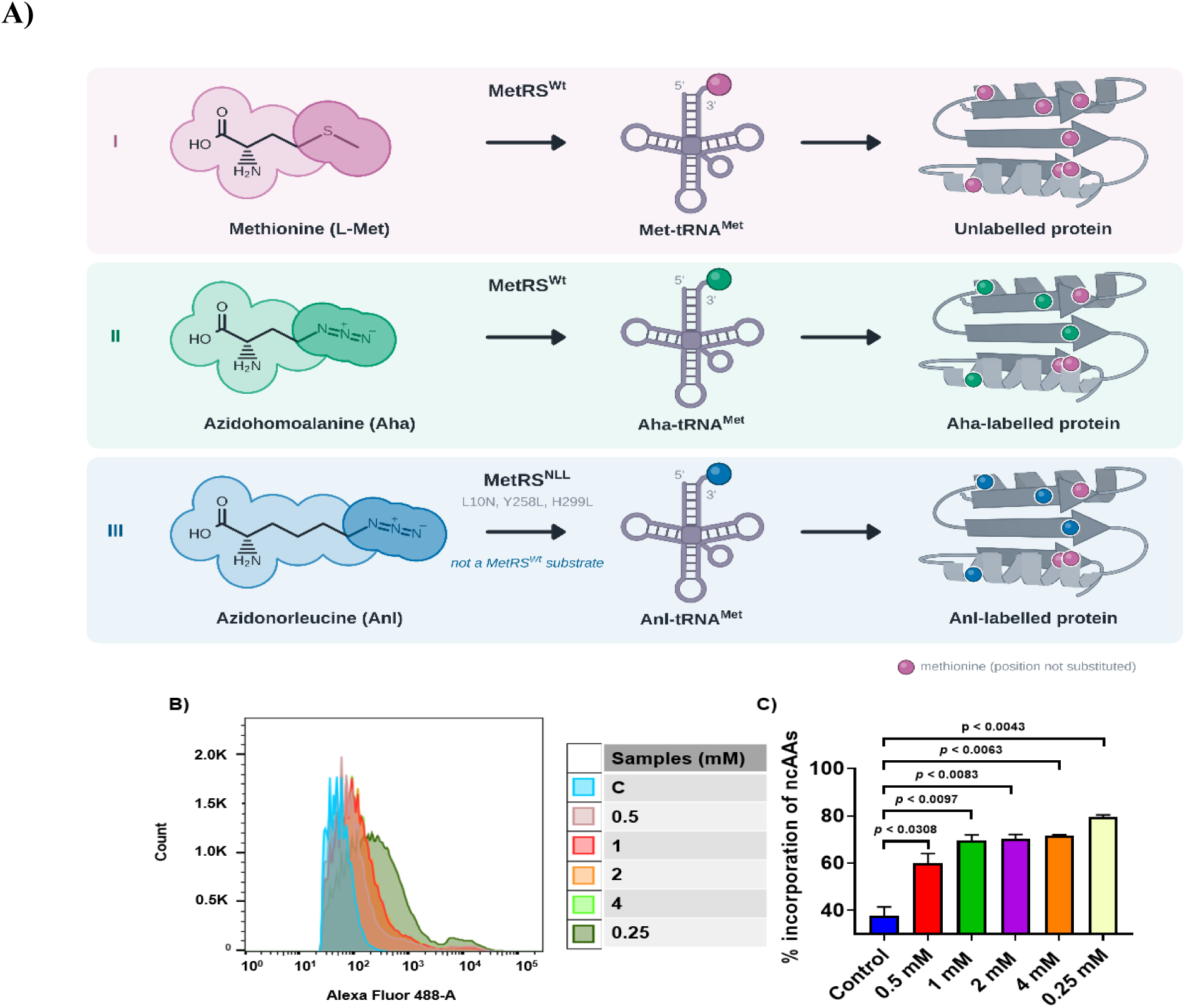
Native and engineered MetRSᴺᴸᴸ of Ab enable metabolic labelling of newly synthesised bacterial proteins using BONCAT. **(A):** Schematic representation of metabolic labelling mediated by native (WT) and engineered MetRSᴺᴸᴸ. Native MetRS charges its cognate tRNA with L-methionine (L-Met; **I**) or the non-canonical methionine analogue L-azidohomoalanine (AHA; **II**), enabling their incorporation into newly synthesized polypeptide chains. In contrast, the engineered MetRSᴺᴸᴸ variant specifically recognizes and charges L-azidonorleucine (ANL; **III**), facilitating its incorporation into nascent bacterial proteins. Since Ab7 expressing MetRSᴺᴸᴸ retains endogenous WT MetRS, both enzymes contribute to protein synthesis, and hence AHA or ANL incorporation does not result in complete replacement of methionine residues in newly synthesized polypeptides (**II and III**). **(B):** Representative flow cytometry histograms showing metabolic labelling of Ab7 co-expressing WT MetRS and MetRSᴺᴸᴸ. Bacterial cultures were induced with 2 % arabinose for 12 h and then pulse-labelled for 4 h without a non-canonical amino acid (control, C) or in the presence of AHA (0.25 mM) or increasing concentrations of ANL (0.5, 1, 2, and 4 mM). Following labelling, incorporated azide groups were conjugated to Alexa Fluor 488–DIBO alkyne through strain-promoted azide–alkyne cycloaddition (SPAAC). Fluorescence was quantified by flow cytometry, with approximately 100,000 bacterial events acquired per sample to determine the proportion of fluorescently labelled cells. **(C):** Quantitative comparison of the percentages of fluorescence-positive Ab7 cells following metabolic labelling with AHA (0.25 mM) or ANL (0.5, 1, 2, and 4 mM). Statistical comparisons were performed using Student’s *t*-test.

To determine whether a corresponding MetRSᴺᴸᴸ variant could be engineered in Ab, we first compared the amino acid sequences of Ab and *E. coli* MetRS using ClustalW. Sequence alignment revealed 57.9 % amino acid identity between the two enzymes (Supplementary Figure 1). Despite this limited sequence identity, structural alignment of the available crystal structures of Ab MetRS (PDB: 5URB) and *E. coli* MetRS (PDB: 1PG2), performed using WinCoot (49), revealed high conservation of the three residues constituting the L-Met-binding pocket that was targeted for mutagenesis: L10, Y258, and H299 in Ab, corresponding to L13, Y260, and H301 in *E. coli* (Supplementary Figure 2). The *E. coli* MetRS structure (PDB: 1PG2) contains an N-terminal His-tag and additional sequence spanning residues 1–21, which was accounted for when comparing residue positions. Together, these sequence and structural analyses identified the corresponding residues in Ab MetRS and provided the rationale for introducing the L10N, Y258L, and H299L substitutions to generate the Ab MetRSᴺᴸᴸ variant. Using site-directed mutagenesis, we generated the Ab MetRSᴺᴸᴸ variant, placed it under the control of the arabinose-inducible pBAD promoter (induced with 2 % arabinose), and introduced the resulting construct into the Ab isolate, Ab7 used in this study (see Materials and Methods for construction details; Supplementary Figure 3).

### The engineered Ab MetRSᴺᴸᴸ variant efficiently incorporates ANL into newly synthesized bacterial proteins

To (i) evaluate the efficiency of ANL incorporation into newly synthesized Ab proteins and (ii) determine an optimal ANL concentration for bio-orthogonal non-canonical amino acid tagging (BONCAT), mid-logarithmic-phase cultures of Ab7 expressing MetRSᴺᴸᴸ were incubated in M9 minimal medium supplemented with increasing concentrations of ANL (0.5, 1, 2, and 4 mM). Labelling efficiency was compared with that achieved using 0.25 mM AHA, a methionine analogue that can be incorporated by endogenous MetRS. Following metabolic labelling, the incorporated azide groups were conjugated to Alexa Fluor 488–DIBO alkyne through strain-promoted azide–alkyne cycloaddition (SPAAC), and fluorescence quantified by flow cytometry (∼100,000 events per sample). Flow cytometric analysis revealed comparable labelling efficiencies at ANL concentrations of 1, 2, and 4 mM (Figure 1B), indicating that increasing ANL concentrations beyond 1 mM did not appreciably improve labelling under the conditions tested. Approximately 70% of Ab7 cells expressing MetRSᴺᴸᴸ were fluorescently labelled following ANL supplementation, compared with approximately 80% of cells labelled with AHA (Figure 1B). The relatively higher proportion of AHA-labelled cells is consistent with the ability of endogenous wild-type MetRS to recognize and incorporate AHA as a methionine analogue (31). Together, these findings demonstrate efficient ANL-dependent metabolic labelling by the engineered Ab MetRSᴺᴸᴸ variant and support its application for selective labelling of newly synthesised bacterial proteins.

### Incorporation of non-canonical amino acids does not compromise Ab7 growth or its viability

Because L-methionine (L-Met) is an essential amino acid for protein function, its substitution with non-canonical amino acid (ncAA) analogues, such as L-azidohomoalanine (AHA) and L-azidonorleucine (ANL), could potentially affect protein functions and consequently bacterial growth and fitness. To assess this possibility, we examined the growth kinetics of Ab7 in M9 minimal medium supplemented either with 0.25 mM AHA or 1 mM ANL for 18 h, with CFU determined at 0, 6, 12 and 18 h. Ab7 expressing endogenous wild-type MetRS was cultured in the presence of AHA, whereas Ab7 co-expressing wild-type MetRS and the engineered MetRSᴺᴸᴸ variant was cultured with ANL. Corresponding cultures grown without ncAA supplementation served as controls.

AHA is a widely used nCAA analogue whose structural similarity to L-Met facilitates its incorporation into newly synthesized proteins with relatively limited perturbation of protein structure and function. (29,31,50), Furthermore, proteomic and metabolic-labelling studies have reported that AHA do not appreciably alter global protein synthesis or degradation in certain bacterial and mammalian cell systems at low concentrations (51,52). Consistent with these observations, our growth kinetics analyses revealed no detectable impairment of Ab7 growth following supplementation with either AHA or ANL under the experimental conditions tested (Supplementary Figure 4). These findings indicate that both ncAAs are compatible with Ab7 growth and support their use for metabolic labelling in subsequent BONCAT experiments.

### L-Methionine biosynthesis inhibitory amino acids reduce endogenous L-methionine levels in Ab7 without compromising bacterial growth

Efficient metabolic incorporation of non-canonical methionine analogues, such as AHA and ANL, requires minimizing competition from endogenous L-methionine (L-Met). Although the engineered MetRSᴺᴸᴸ variant recognizes ANL, the endogenous wild-type MetRS incorporates L-Met for protein synthesis at higher efficiency (30–32,48). Because L-Met is essential for bacterial growth and survival, its intracellular availability must be reduced without compromising bacterial fitness. Accordingly, we investigated the effects of methionine biosynthesis inhibitory amino acids (MBIAA) and sodium acetate (SA) on endogenous L-Met levels in Ab7. Previous studies have demonstrated that MBIAA can facilitate metabolic labelling with AHA in *E. coli* by suppressing endogenous methionine biosynthesis (39). Similarly, exposure of E. coli to 8 mM acetate at pH 6.0 has been reported to impair methionine biosynthesis and promote intracellular homocysteine accumulation, contributing to growth inhibition (53).

To determine the effectiveness of these approaches in Ab7, we performed metabolomic analyses of WT Ab7 cells grown in M9 media following treatment with MBIAA or SA. MBIAA treatment significantly reduced intracellular L-Met levels by 16 h compared with SA treatment (Fig. 2A), without appreciably affecting bacterial growth kinetics (Supplementary Figure 5). Principal component analysis (PCA) of the metabolomic profiles at the respective sampling time points is presented in Supplementary Figure 6. These findings indicate that MBIAA treatment effectively reduces endogenous L-Met availability while preserving Ab7 growth, supporting its use to facilitate metabolic incorporation of L-methionine analogues in subsequent BONCAT experiments. SA at 8 mM neither did exhibit any inhibitory effect on Ab7 growth nor significantly reduced L-Met levels (Fig. 2A).

**Figure 2.**
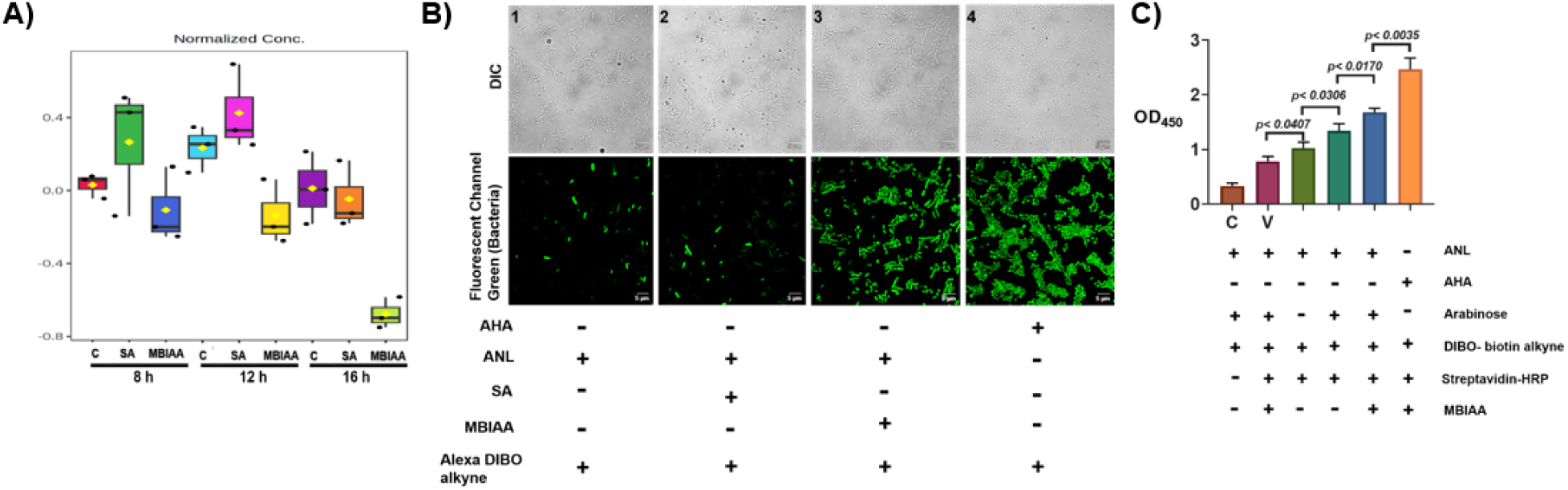
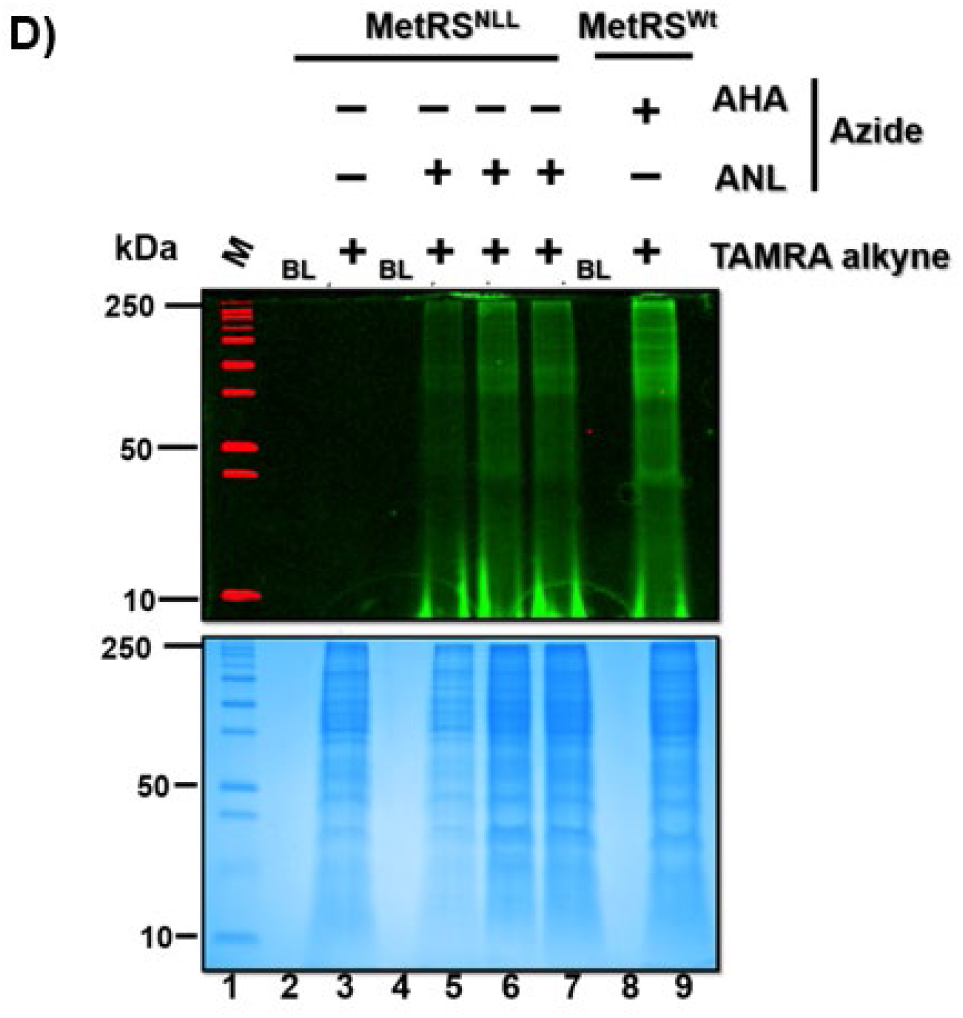
Methionine biosynthesis inhibitory amino acids (MBIAA) reduce endogenous L-methionine levels and support efficient ANL incorporation into newly synthesized Ab proteins. **(A):** Effect of MBIAA and sodium acetate (SA) on intracellular L-methionine (L-Met) levels in Ab7. Box plots represent the relative abundance (fold change) of L-Met in untreated control, SA-treated, and MBIAA-treated cultures at 8, 12, and 16 h. Statistical significance was assessed by one-way analysis of variance (ANOVA) using MetaboAnalyst (*P* < 0.05). Group colors were assigned automatically by the software. Data in (A) were obtained from three biological replicates. **(B):** Representative confocal fluorescence microscopy images (top - DIC and bottom - green fluorescent channel) showing incorporation of non-canonical amino acids (ncAAs) into Ab7 proteins. Following metabolic labelling, bacterial cells were labelled with 15 µM Alexa Fluor 488–DIBO alkyne through strain-promoted azide–alkyne cycloaddition (SPAAC) to fluorescently label incorporated azide groups. Images were acquired at 60X magnification. Green fluorescence indicates ncAA-labelled Ab7 cells. Scale bars, 5 µm (white). **(C):** Quantitative assessment of ANL incorporation into Ab7 proteins expressing MetRSᴺᴸᴸ in the presence of MBIAA using a biotin–streptavidin-based detection assay. Following metabolic labelling, incorporated azide groups were conjugated to 20 µM DIBO–biotin alkyne through SPAAC. Biotinylated proteins were subsequently detected using streptavidin–horseradish peroxidase (HRP), with 3,3′,5,5′-tetramethylbenzidine (TMB) as the chromogenic substrate. Statistical significance was determined using Student’s *t*-test. Exact *P* values for the indicated comparisons were 0.0407, 0.0306, 0.0170, and 0.0035 (*P* < 0.05 for the first three comparisons; *P* < 0.01 for the fourth). **(D):** Comparative analysis of ANL incorporation mediated by engineered MetRSᴺᴸᴸ and AHA incorporation mediated by endogenous wild-type MetRS. Following metabolic labelling and conjugation of incorporated azide groups to TAMRA–alkyne, proteins were resolved by SDS–PAGE. In-gel fluorescence was detected using an Azure Biosystems c400 gel imager in the Cy3 channel (excitation, 526 nm; emission, 565 nm). The same gel was subsequently stained with Coomassie Brilliant Blue to assess protein loading. M, molecular weight marker with representative protein sizes in kDa; BL, blank.

### MBIAA supplementation enhances ANL incorporation into newly synthesized Ab proteins by **MetRSᴺᴸᴸ**

To evaluate the enhancement of ANL incorporation into newly synthesized Ab proteins in the presence of methionine biosynthesis inhibitory amino acids (MBIAA), we employed three complementary approaches: confocal fluorescence microscopy, biotin–streptavidin-based detection, and in-gel fluorescence analysis. Confocal microscopy and biotin–streptavidin assays demonstrated efficient ANL incorporation by the engineered MetRSᴺᴸᴸ variant in the presence of MBIAA and 1 mM ANL (Fig. 2B and 2C). Fluorescence intensity was quantified using corrected total cell fluorescence (CTCF) measurements (Supplementary Figure 7). Furthermore, in-gel fluorescence analysis following copper-catalyzed azide–alkyne cycloaddition (CuAAC) with TAMRA–alkyne revealed ANL-dependent protein labelling comparable to that achieved with AHA incorporation mediated by endogenous wild-type MetRS (Fig. 2D). Collectively, these findings confirm that the engineered Ab MetRSᴺᴸᴸ variant efficiently incorporates ANL into newly synthesized bacterial proteins under MBIAA-supplemented conditions, supporting its suitability for subsequent BONCAT-based proteomic analyses.

### Ab7 persists within A549 lung epithelial cells to a comparable extent to other clinical isolates

To evaluate the intracellular survival of Ab in human lung epithelial cell line A549, we determined bacterial colony-forming units (CFUs) following infection of A549 cells with Ab7 and two additional isolates: Ab755 (from our laboratory collection) and AB5075, originally isolated from a patient with tibial osteomyelitis (54). CFU enumeration demonstrated that Ab7 persisted within A549 cells, with intracellular counts comparable to those of Ab755 and AB5075 (Supplementary Figure 8). These findings demonstrate that all three isolates persist within A549 epithelial cells to a comparable extent.

### Metabolic labelling with AHA does not compromise the intracellular survival of Ab7 in A549 epithelial cells

To determine whether metabolic labelling with non-canonical amino acids (ncAAs) affects the intracellular survival of Ab7, we compared the infection kinetics of AHA-labelled Ab7 expressing MetRSᴺᴸᴸ with those of unlabelled wild-type Ab7 in A549 lung epithelial cells. Briefly, Ab7 expressing MetRSᴺᴸᴸ was initially cultured in LB medium and subsequently subcultured into M9 minimal medium at an initial OD₆₀₀ of 0.005; at OD₆₀₀ 0.1, 0.25 mM AHA and MBIAA were added and metRSᴺᴸᴸ was induced with 2 % arabinose for 12 h. Following metabolic labelling, bacteria were pelleted and washed three times with PBS (pH 7.4) to remove residual extracellular AHA. A549 cells were then infected at a multiplicity of infection (MOI) of 1:1000 for 3 h in L-methionine-depleted DMEM. Following elimination of extracellular bacteria, intracellular bacterial survival was assessed by CFU enumeration at different post-infection time points. AHA-labelled Ab7 exhibited intracellular survival kinetics comparable to those of unlabelled wild-type Ab7, indicating that metabolic labelling did not appreciably compromise bacterial survival (for the duration tested) within A549 cells (Supplementary Figure 9).

### MetRSᴺᴸᴸ enables ANL incorporation into newly synthesized Ab7 proteins during infection of A549 epithelial cells

To confirm ANL incorporation into newly synthesized Ab7 proteins during infection of human lung epithelial cells, A549 cells were infected with Ab7 expressing MetRSᴺᴸᴸ at a multiplicity of infection (MOI) of 1:1000 and incubated for 3 h in L-methionine-depleted DMEM supplemented with 1 mM ANL. Following infection, cells were washed with PBS (pH 7.4), permeabilised, and subjected to strain-promoted azide–alkyne cycloaddition (SPAAC) using Alexa Fluor 488–DIBO alkyne to fluorescently label ANL-containing proteins. Confocal fluorescence microscopy revealed labelled bacteria within A549 cells, demonstrating successful ANL incorporation into bacterial proteins during infection (Fig. 3A). To determine whether ANL was incorporated into bacterial proteins across a broad range of molecular weights, total lysates from infected A549 cells were subjected to copper-catalyzed azide–alkyne cycloaddition (CuAAC) using tetramethylrhodamine (TAMRA)–alkyne, followed by in-gel fluorescence analysis (Fig. 3B). Fluorescently labelled proteins were detected across a wide molecular weight range (10 to 250 kDa), demonstrating that ANL incorporation was not restricted to proteins of a particular size. Together, confocal microscopy and in-gel fluorescence analyses confirmed efficient ANL incorporation into a broad repertoire of newly synthesized Ab7 proteins during infection of A549 epithelial cells.

**Figure 3.**
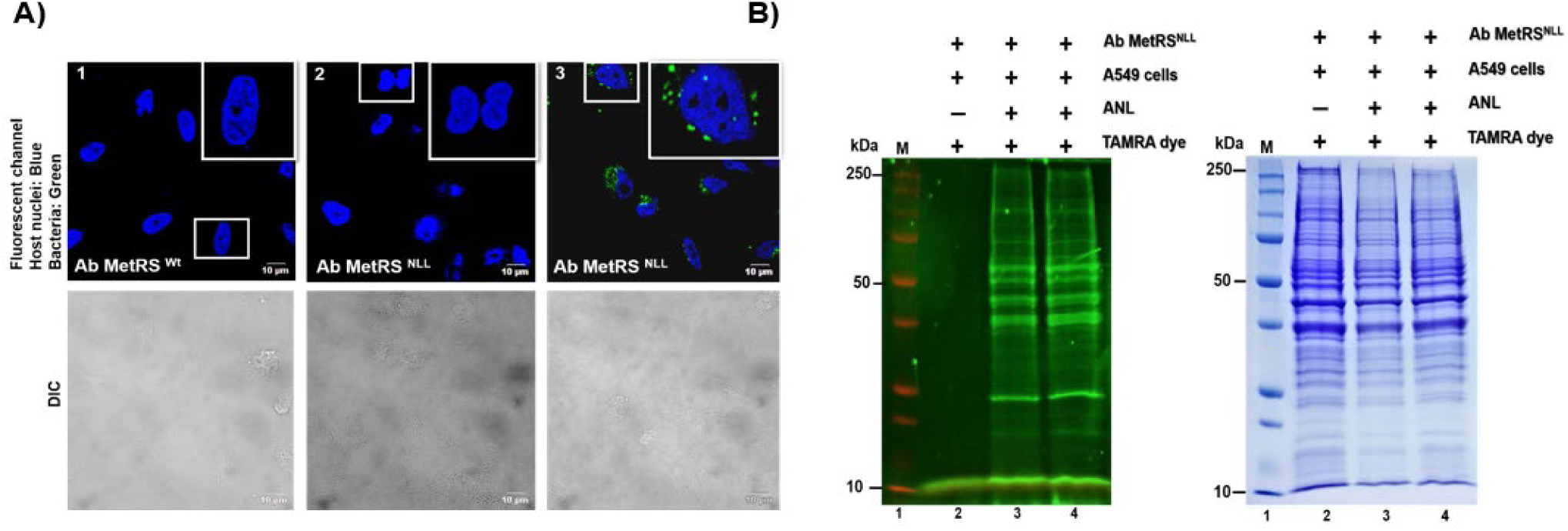
In situ detection of ANL-labelled Ab and its proteins during infection of A549 lung epithelial cells. **(A):** Confocal fluorescence microscopy demonstrating ANL incorporation into Ab7 proteins during infection of A549 lung epithelial cells. A549 cells were infected with Ab7 expressing MetRSᴺᴸᴸ at a multiplicity of infection (MOI) of 1:1000 in the presence of 1 mM ANL. Following infection, cells were fixed and permeabilized. Host-cell nuclei were stained with 4′,6-diamidino-2-phenylindole (DAPI; blue), while ANL-labelled bacterial proteins were detected using Alexa Fluor 488–DIBO alkyne (green) through strain-promoted azide–alkyne cycloaddition (SPAAC). Fluorescence signals were visualized by confocal microscopy. Scale bar, 10 µm. **(B):** In-gel fluorescence analysis of ANL-labelled Ab7 proteins recovered from infected A549 cells. Following infection, total cell lysates were subjected to copper-catalyzed azide–alkyne cycloaddition (CuAAC) with tetramethylrhodamine (TAMRA)–alkyne. Proteins were subsequently precipitated using methanol–chloroform to remove unreacted TAMRA–alkyne before SDS–PAGE and fluorescence imaging. Three biological replicates were performed; two are shown here. Lane 1, molecular weight marker (M); lane 2, A549 cells infected with Ab7 in the absence of ANL (negative control); lanes 3 and 4, A549 cells infected with Ab7 in the presence of ANL (biological triplicates). The same gel was subsequently stained with Coomassie Brilliant Blue to assess protein loading.

### BONCAT and conventional proteomics enable identification of the infection-associated Ab protein in A549 epithelial cells

Our in-gel fluorescence analysis demonstrated the incorporation of ANL into newly synthesised Ab7 proteins during infection of A549 epithelial cells (Fig. 3B). However, this approach did not establish the identities of the labelled proteins or distinguish proteins synthesised before infection from those synthesised during infection. To address these questions, we employed a dual metabolic-labeling strategy using two non-canonical amino acids (ncAAs), AHA and ANL, in combination with conventional proteomics (Fig. 4A).

**Figure 4.**
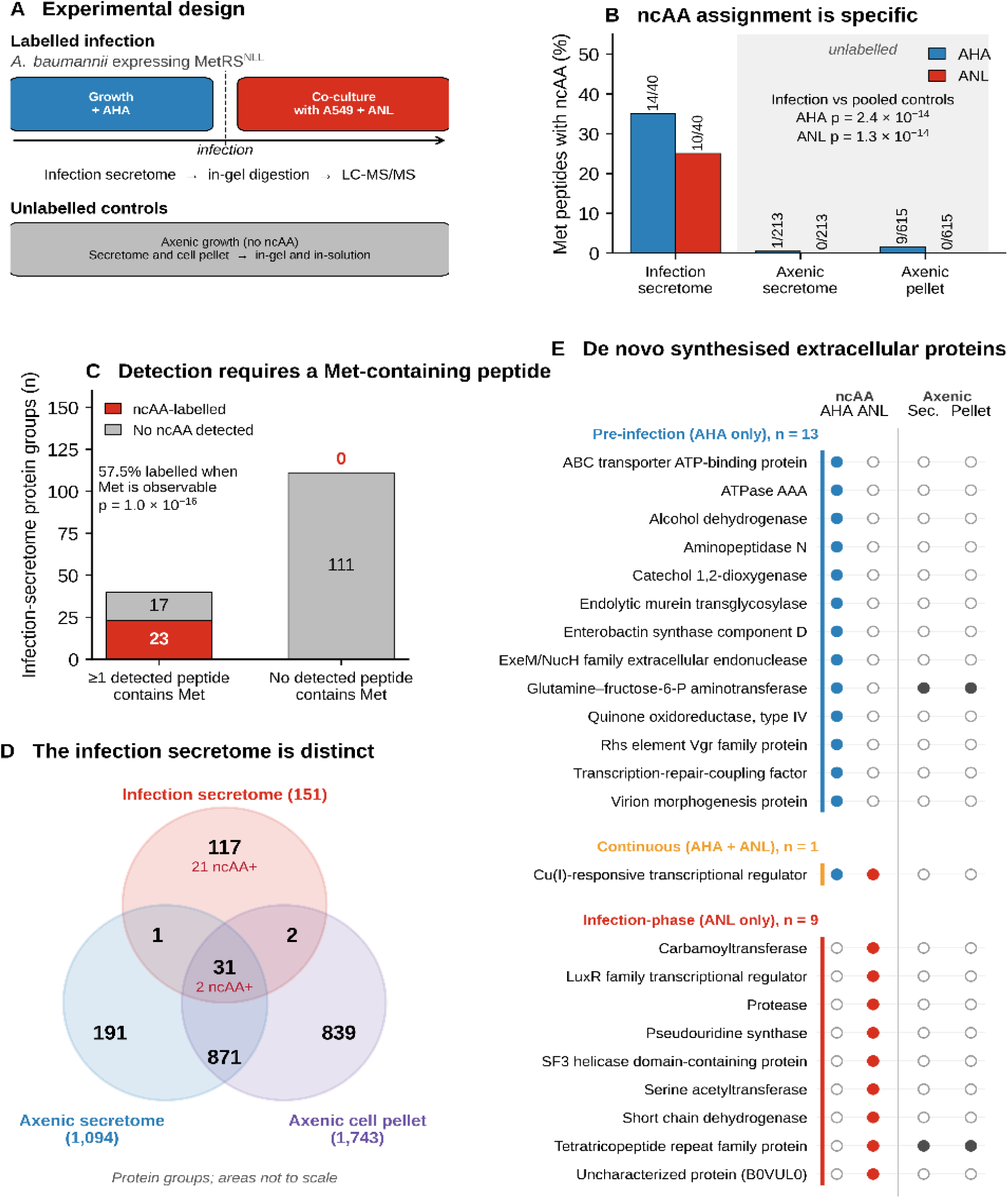
Sequential non-canonical amino acid labelling distinguishes Ab proteins synthesised before and during infection of A549 cells. **(A):** Experimental design. Ab co-expressing the engineered methionyl-tRNA synthetase variant MetRSᴺᴸᴸ was grown for 12 h in the presence of 0.25 mM AHA, followed by co-culture with A549 cells in medium containing 1 mM azidonorleucine ANL. The cell-free infection-associated Ab protein was subjected to in-gel tryptic digestion and LC–MS/MS analysis. Unlabelled axenic culture-filtrate proteome and axenic bacterial cell lysate samples, grown without either analogue, served as negative controls. **(B):** Detection of AHA- and ANL-modified methionine-containing peptides in the infection-associated Ab protein compared with two unlabelled axenic in-gel datasets (828 peptides combined). Numbers above the bars indicate modified peptides relative to the total number of methionine-containing peptides. Statistical significance was determined using a two-sided Fisher’s exact test against the pooled controls. **(C):** Classification of the 151 infection associated Ab protein protein groups according to the presence or absence of methionine-containing peptides among those detected, with the number of protein groups containing site-localised analogue modifications indicated. The 111 protein groups shown on the right contain methionine residues in their full-length sequences but lack methionine-containing peptides among those identified, precluding direct detection of analogue incorporation. Statistical significance was assessed using a two-sided Fisher’s exact test. **(D):** Overlap of identified protein groups across the three experimental conditions. Axenic datasets combine identifications from in-gel and in-solution digestion. Red numbers indicate analogue-labelled protein groups. Circled areas are not proportional to the number of protein groups. **(E):** Classification of the 23 protein groups containing site-localized AHA- or ANL-modified peptides according to their respective synthesis windows. Filled circles under the ncAA columns indicate detection of AHA- or ANL-modified peptides, whereas filled circles under the axenic columns indicate detection of the corresponding protein group in the protein (Sec.) or axenic bacterial cell lysate. Data from three biological replicates were searched independently, and the identification results were merged. Peptide and protein identifications were accepted at a false discovery rate <1%.

Ab7, containing endogenous wild-type MetRS, engineered to also express MetRSᴺᴸᴸ variant, was grown in M9 minimal medium supplemented with 0.4% succinate (at OD₆₀₀ 0.005), 2% arabinose and MBIAA (at OD₆₀₀ ∼0.1). Pre-infection metabolic labelling was achieved for 12 h with AHA (0.25 mM). AHA was incorporated by endogenous wild-type MetRS. Following this, bacteria were harvested, washed with 1X PBS (pH 7.4), and resuspended in L-methionine-free DMEM for infection of A549 cells.

AHA labelled Ab7, also expressing the MetRSᴺᴸᴸ variant, was used at MOI 1:1000 to infect 10⁵ A549 cells in a L-methionine-free DMEM supplemented with 1 mM ANL. Unlike AHA, ANL gets incorporated by only the engineered MetRSᴺᴸᴸ variant. After 3 h of infection, the monolayers were washed and treated with colistin (8 µg/mL) for 1 h to eliminate extracellular bacteria. The cells were subsequently washed and incubated for an additional 9 h in fresh ANL-containing medium, completing the infection-associated labelling period.

Following infection, A549 cells were lysed using chilled water supplemented with protease inhibitors. The resulting lysates were centrifuged and passed through 0.2-µm filters to remove intact bacteria. The recovered cell-free protein fractions were separated by SDS–PAGE, subjected to in-gel tryptic digestion, and analyzed by LC–MS/MS using a TripleTOF 5600 mass spectrometer. Protein identifications were accepted at a false discovery rate (FDR) of 1%. Unlabelled axenic bacterial culture filtrate and axenic bacterial cell lysate were processed in parallel to generate reference proteomes.

Together, BONCAT and conventional proteomics were enabled as complementary approaches to characterize the infection-associated Ab7 secreted protein repertoire that was recovered from the A549 cellular milieu. Sequential labelling with AHA and ANL distinguished bacterial proteins synthesised before infection from those synthesised during infection, including proteins detected in both labelling windows (Fig. 4). Conventional proteomics additionally enabled the identification of bacterial proteins irrespective of analogue incorporation, thereby expanding coverage beyond the metabolically labelled proteome.

### Sequential L-Met analogue labelling distinguishes two translational windows

AHA present during pre-infection is incorporated into bacterial proteins by endogenous wild-type methionyl-tRNA synthetase (MetRS), whereas ANL present during co-culture with A549 cells is selectively incorporated by the engineered MetRSᴺᴸᴸ variant (Fig. 4A). Thus, AHA-modified peptides primarily represented proteins synthesized during pre-infection labelling, although residual intracellular AHA following washing could contribute to labelling during co-culture. In contrast, ANL-modified peptides provided evidence of bacterial protein synthesis during co-culture. Importantly, detection of either analogue alone does not establish that the corresponding protein was synthesized exclusively within that labelling period. Nevertheless, all the labelled proteins (with AHA or ANL alone or combined) were detected from the host cellular milieu indicating possible release of AHA labelled Ab proteins into host cellular milieu.

Following the exclusions described in Methods, LC–MS/MS analysis of the infection-associated host cellular milieu fraction identified 179 peptides corresponding to 152 Ab proteins (Supplementary Table S3). Throughout this study, proteins are reported as protein groups, because few accessions share an identical peptide (of at least seven amino acid residues) and they could not be distinguished using the available peptide evidence (Supplementary Table S4). Hence, of these 152 Ab protein groups, 23 protein groups contained at least one peptide with a site-localized AHA or ANL modification (Supplementary Table S5) and the summary of labelling efficiency is detailed in Supplementary Table S6. Contaminant human proteins are as listed in Supplementary Table S7.

### Specificity of amino acid analogue assignment

Unlabeled axenic samples, analyzed using identical database search parameters, were used to estimate background analogue assignments. The control dataset comprised two in-gel digestion datasets containing 213 and 615 methionine-containing peptides, respectively (828 peptides in total; in-solution datasets were excluded as described in Methods; Supplementary Table S8). Among these, AHA modifications were assigned to 10 peptides (1.21%; 95% CI: 0.58–2.21%), whereas no ANL modifications were detected (0%; upper 95% confidence bound: 0.44%). In contrast, among the 40 methionine-containing peptides identified in the infection-associated Ab protein fraction, 14 carried AHA modifications and 10 carried ANL modifications (Fig. 4B; Supplementary Table S9). Both analogue assignments were significantly enriched relative to the unlabeled controls (AHA, p = 2.4 × 10⁻¹⁴; ANL, p = 1.3 × 10⁻¹⁴). Because efficient ANL incorporation requires the engineered MetRSᴺᴸᴸ variant and ANL was supplied exclusively during co-culture, ANL-modified peptides provide direct evidence of bacterial protein synthesis during host-cell infection.

### Detection of analogue incorporation is constrained by methionine-containing peptides

Analogue incorporation can be detected only in peptides containing methionine residues. Of the 179 peptides identified in the infection-associated Ab protein fraction, 40 contained methionine, of which 23 (57.5%) carried an AHA or ANL modification. At the residue level, 24 of 49 methionine positions (49.0%) were substituted with an analogue (Supplementary Table S3 – S6). As expected, none of the 139 methionine-free peptides carried an analogue modification. At the protein-group level, 23 of the 40 groups identified through at least one methionine-containing peptide were labelled, compared with none of the 111 groups identified exclusively through methionine-free peptides (Fig. 4C; p = 1.0 × 10⁻¹⁶). Thus, the detectable labelled proteome is constrained primarily by the recovery of methionine-containing peptides, rather than simply by analogue incorporation efficiency. Importantly, the absence of a detectable analogue modification does not establish that a protein was not synthesised during the labelling period.

### The infection-associated Ab protein secretome repertoire differs substantially from the axenic culture filtrate proteome

The infection-associated Ab protein repertoire comprised 151 protein groups, compared with 1,094 identified in the axenic culture-filtrate proteome and 1,743 in the axenic bacterial cell pellet (Supplementary Table S10, S11). Notably, 117 of the 151 infection-associated protein groups (77.5%) were not detected in either axenic dataset, including 21 of the 23 analogue-labelled protein groups (Figure 4D; Supplementary Table S12). These findings reveal a substantial difference between the proteins recovered during infection and those detected under axenic conditions. However, because the axenic datasets provided greater proteomic coverage, these differences should be interpreted as differences in detection rather than definitive evidence that the corresponding proteins are absent under axenic conditions.

### The infection-associated Ab secreted protein fraction is enriched in predicted secretory and outer-membrane proteins

Compared with protein groups detected exclusively in axenic bacterial cell pellet, the infection-associated Ab secreted protein fraction showed significant enrichment for proteins containing predicted signal peptides (25/151, 16.6% versus 49/841, 5.8%; odds ratio [OR] = 3.21, p = 2.9 × 10⁻⁵; Supplementary Figure 11C, Supplementary Tables S13 and S14) and proteins predicted to localize to the outer membrane (8/151, 5.3% versus 12/841, 1.4%; OR = 3.86, p = 0.006; Supplementary Figure 11D, Supplementary Tables S15 and S16). Nevertheless, cytoplasmic proteins constituted the largest predicted localization category across all fractions, including the axenic culture-filtrate proteome. The enrichment of signal peptide-containing and outer-membrane proteins is consistent with preferential recovery of exported or envelope-associated bacterial proteins during infection, although contributions from bacterial lysis and outer membrane extracellular vesicles of Ab cannot be excluded (Supplementary Table S17).

### Functional annotation of infection-associated Ab secreted proteins

Among the 117 protein groups detected exclusively in the infection-associated fraction, 46 had KEGG pathway annotations (Supplementary Table S18). Of the 19 pathways meeting the testing threshold, two were significantly enriched relative to the background of 957 annotated protein groups: bacterial secretion system (6/46 versus 31/957; 4.0-fold enrichment, q = 0.025) and exopolysaccharide biosynthesis (3/46 versus 6/957; 10.4-fold enrichment, q = 0.025; Supplementary Figure 12,

Supplementary Table S18). Proteins contributing to the bacterial secretion system pathway included a VgrG-family type VI secretion system protein, the protein translocase subunit SecD, a TolC-family protein, and a type II secretion system protein L. Notably, the VgrG-family protein contained an AHA modification, indicating synthesis during the pre-infection labelling period. In contrast, enrichment of the exopolysaccharide biosynthesis pathway was driven largely by two serine acetyltransferases assigned to this pathway by KEGG based on secondary enzymatic activities; consequently, this enrichment was not considered further.

No KEGG pathway was significantly enriched when the entire infection-associated Ab protein repertoire was analyzed. Although Pfam domains were identified in 124 of the 151 protein groups, no domain occurred in three or more groups, indicating considerable functional heterogeneity (Supplementary Table S12). Gene Ontology (GO) enrichment analysis was not performed because eggNOG-mapper transferred annotations from assigned orthologous groups, including eukaryote-specific GO terms that were inappropriate for these bacterial proteins (Supplementary Table S19). Functional interpretation was therefore restricted to KEGG pathways and sequence-derived domain annotations.

### Labelling profiles of infection-associated Ab secreted proteins

The 23 analogue-labelled protein groups were classified as AHA-only, ANL-only, or dual-labelled based on their detected modifications (Figure 4E; Supplementary Table S5, S12). ANL incorporation provided evidence of bacterial protein synthesis during co-culture with A549 cells, whereas AHA incorporation was primarily associated with the pre-infection labelling period, although residual intracellular AHA could contribute to labelling during co-culture. Thus, these categories reflect the modifications detected rather than exclusive synthesis within a particular labelling window.

Thirteen protein groups were identified exclusively through AHA-modified peptides: an ABC transporter ATP-binding protein, AAA-family ATPase, alcohol dehydrogenase, aminopeptidase N, catechol 1,2-dioxygenase, endolytic murein transglycosylase, enterobactin synthase component D, ExeM/NucH-family extracellular endonuclease, glutamine-fructose-6-phosphate aminotransferase, type IV quinone oxidoreductase, Rhs element Vgr-family protein, transcription-repair coupling factor, and virion morphogenesis protein. Nine protein groups were identified exclusively through ANL-modified peptides, supporting their synthesis during co-culture: a carbamoyltransferase, LuxR-family transcriptional regulator, protease, pseudouridine synthase, SF3 helicase domain-containing protein, serine acetyltransferase, short-chain dehydrogenase, tetratricopeptide repeat-family protein, and an uncharacterized protein (B0VUL0; Supplementary Table S3).

The remaining protein group, a Cu(I)-responsive transcriptional regulator, was identified through a peptide containing both AHA and ANL modifications at distinct methionine residues (MNIGKVSQASGVTAKMIR; AHA at Met1 and ANL at Met16; Supplementary Table S3). Assuming accurate modification assignment, the presence of ANL indicates synthesis of the modified peptide during co-culture. However, the simultaneous detection of both analogues within the same peptide does not establish that the protein was synthesized in both labeling periods. Rather, it is consistent with incorporation of residual intracellular AHA alongside ANL during co-culture.

## Discussion

Ab, a major ESKAPE pathogen, causes pneumonia, bloodstream infections, and sepsis, posing a substantial clinical challenge, particularly in low- and middle-income countries. Etiological diagnosis remains difficult in neonatal sepsis, where limited blood volumes and the low sensitivity of blood cultures can hinder pathogen identification. We hypothesised that bacterial proteins released during infection could provide alternative targets for pathogen-specific diagnosis, including in culture-negative infections. With the long-term objective of developing monoclonal antibody- or aptamer-based assays to detect such proteins, we sought to characterise the repertoire of infection-associated secreted proteins of Ab. Using the human lung epithelial cell line A549 as an infection model, we combined BONCAT with conventional proteomics, recognizing that neither approach alone could provide sufficiently comprehensive coverage of the Ab proteins released into the host-cell environment during infection. The complementary value of these approaches is particularly important because metabolic labelling establishes when bacterial proteins are synthesised, whereas conventional proteomics can recover proteins for which analogue incorporation is not detectable. Their combined application therefore addresses both the temporal origin and breadth of the infection-associated bacterial protein repertoire.

Identifying Ab-derived proteins released during infection requires distinguishing the bacterial proteome from the abundant host-cell proteome. Although BONCAT offers a means of selectively labelling newly synthesised proteins, its application to host–pathogen interactions is complicated by the high substrate specificity of aminoacyl-tRNA synthetases (aaRSs), which generally restricts and prevent any incorporation of non-canonical amino acids (ncAAs). Interestingly, AHA, a very close structural analogue of L-methionine (L-Met), can nevertheless be recognized by endogenous methionyl-tRNA synthetases (MetRSs) and incorporated into newly synthesized proteins. However, because of structural conservation across kingdoms, this recognition occurs in both bacterial and mammalian cells, and thus AHA cannot be exploited to selectively distinguish *Ab*-derived proteins from those synthesized by the host. Consequently, engineering a bacterial MetRS capable of incorporating an orthogonal ncAA became a necessity for selective labelling of pathogen-derived proteins during infection.

To achieve this selectivity, we exploited ANL, another structural analogue of L-Methionine that bears a terminal azide on an extended side chain making it ineffective in getting accommodated into the substrate-binding pocket of wild-type methionyl-tRNA synthetase (MetRS^LYH^). Unlike AHA, ANL requires an engineered MetRS for efficient incorporation into newly synthesized proteins. Tanrikulu et al. (2009) (48), demonstrated that *E. coli* MetRS variants differ substantially in their substrate preferences: L13G, CLL, and SLL variants preferentially activate L-methionine and exhibit poor ANL activation, whereas PLL and NLL variants efficiently activate ANL while discriminating against L-methionine, enabling selective protein labelling under physiological amino acid concentrations. Given the subsequent application of the NLL variant in bacterial and mammalian cell models, we engineered the corresponding variant in Ab by introducing three substitutions (L10N, Y258L, and H299L) into the L-methionine-binding pocket, equivalent to L13N, Y260L, and H301L in *E. coli* MetRS.

While engineering MetRSᴺᴸᴸ enabled selective incorporation of ANL into *Ab* proteins, achieving efficient labelling without compromising bacterial physiology presented an additional challenge. Extensive replacement of L-methionine with ANL could potentially alter protein structure and function, thereby affecting bacterial viability, growth, survival, and pathogenicity. We therefore sought to balance sufficient ANL incorporation for proteomic detection with minimal physiological perturbation. Because Ab synthesizes its own L-methionine, limiting its availability required reducing the intracellular pool rather than merely eliminating exogenous methionine. We first optimized growth in minimal medium, identifying 0.4% succinate as the preferred carbon source for Ab7, and subsequently employed M9 minimal medium to restrict exogenous L-methionine. To further reduce intracellular L-methionine availability, we evaluated methionine biosynthesis inhibitory amino acids (MBIAA) and sodium acetate (SA), monitoring intracellular L-methionine levels by metabolomics. While SA produced no measurable reduction, MBIAA significantly decreased intracellular L-methionine levels by 16 h without appreciably affecting bacterial growth. Accordingly, MBIAA supplementation in M9 medium was adopted to facilitate ANL labelling while preserving bacterial growth.

Having optimised ANL incorporation while minimising its effects on bacterial growth, we next verified ncAA labelling using three complementary approaches: confocal microscopy, a biotin–streptavidin interaction assay, and in-gel fluorescence. Confocal microscopy confirmed the presence of ANL-labelled Ab7 within infected A549 cells under methionine-free DMEM conditions. Furthermore, labelled Ab7 persisted intracellularly, exhibiting a pattern comparable to that of the hypervirulent AB5075 strain, while ncAA incorporation did not appreciably alter its growth dynamics under the conditions tested. Together, these observations indicate that ANL labelling was compatible with bacterial growth and intracellular survival, supporting its application to investigate the infection-associated secreted proteome of Ab.

Beyond selective identification of bacterial proteins, our sequential labelling strategy provides an additional advantage: the ability to distinguish proteins synthesized before infection from those produced during host–pathogen interactions. Unlike transcriptomics and conventional steady-state proteomics, which measure transcript abundance and accumulated protein levels, respectively, sequential incorporation of two methionine analogues allows bacterial proteins to be assigned to distinct translational windows. In our approach, AHA was used to label proteins synthesised before infection (in axenic cultures), whereas ANL, incorporated selectively by the engineered MetRSᴺᴸᴸ, marked proteins synthesised during infection. Importantly, the specificity of these assignments was experimentally evaluated rather than assumed. Among 828 methionine-containing peptides identified from bacteria never exposed to either analogue, AHA was assigned to ten peptides, whereas no ANL assignments were detected (Figure 4B). These observations correspond to an apparent background assignment rate of 1.2% for AHA and an approximate upper 95% confidence bound of 0.4% for ANL. Applied to the 40 methionine-containing peptides recovered from the infection-associated extracellular fraction, these rates suggest fewer than one expected background assignment per labelling channel, supporting the specificity of temporal proteins identification.

Nevertheless, the proportion of the newly synthesized proteome accessible through this strategy is constrained by several biological and analytical factors. Proteins differ in their rates of synthesis, abundance, stability, and turnover, and rapidly degraded proteins may escape detection. More fundamentally, identification of an analogue-containing peptide requires the presence of a methionine residue within a peptide detectable after proteolytic digestion. Consequently, proteins identified exclusively through methionine-free peptides cannot be assigned to either labelling window, even if they incorporate an analogue elsewhere in their sequence. Consistent with this limitation, 111 of the 151 identified protein groups were detected only through methionine-free peptides (Fig. 4C). Thus, the 23 labelled proteins represent a detectable subset, rather than the complete repertoire, of proteins synthesised across the two labelling windows. Given the relatively small numbers of proteins assigned to the three labelling categories (13, 1, and 9; Figure 4E), their biological significance was assessed primarily through individual protein identities and known functions rather than statistical enrichment analyses. Importantly, conventional proteomics mitigated this detection limitation: although only 23 of 151 infection-associated protein groups had detectable analogue-modified peptides, the full analysis identified 151 groups, including 111 represented exclusively by methionine-free peptides (Figure 4C; Supplementary Table S4). Thus, BONCAT provided temporal evidence for a defined subset, whereas conventional identification substantially broadened the candidate repertoire without requiring the incorporation of observable analogues.

Despite this limited coverage, several proteins synthesized during co-culture are associated with biological processes implicated in Ab pathogenesis. The LuxR-family regulator is associated with the AbaR/AbaI quorum-sensing system, which regulates biofilm formation, surface-associated motility, and virulence-associated phenotypes (55,56). Serine acetyltransferase participates in cysteine biosynthesis, a pathway implicated in oxidative stress resistance and bacterial fitness during infection (57,58). Notably, a Cu(I)-responsive regulator was the only protein detected in both labelling windows. Copper is an important component of host antimicrobial defence, and Ab possesses copper-responsive regulatory and efflux systems associated with resistance to copper toxicity and virulence (59,60). However, these three proteins are predicted to be cytoplasmic and lack recognizable export signals. Their identification in the extracellular fraction therefore establishes their extracellular recovery but does not demonstrate active secretion or an extracellular function. In contrast, the tetratricopeptide repeat-containing protein was predicted to be a lipoprotein and belongs to a protein family previously implicated in host-cell interactions and interference with phagolysosomal maturation (61). The remaining infection-phase proteins have no established or known roles in Ab pathogenesis. Collectively, these findings identify candidate proteins whose synthesis during host contact warrants further functional investigation.

Beyond identifying proteins synthesized during distinct experimental windows, the integration of BONCAT with conventional proteomics provides insight into how the Ab secreted protein repertoire recovered from infected epithelial cells differs from that observed under axenic conditions. The relative representation of proteins associated with bacterial export pathways and the cell envelope suggests that host-associated conditions may influence the composition of the extracellular bacterial protein repertoire. This is particularly relevant because surface-associated and exported proteins participate in bacterial interactions with host cells and may contribute to processes such as adhesion, envelope remodeling, nutrient acquisition, and adaptation to host-imposed stress. Nevertheless, their extracellular recovery does not necessarily establish active secretion, as proteins may also be released through outer-membrane vesicles, envelope turnover, or bacterial lysis. Importantly, the combined proteomic approach provides a framework for distinguishing proteins with evidence of synthesis during infection from the broader repertoire recovered in the host-associated environment. This distinction may help prioritise candidates for subsequent functional characterisation and evaluation as infection-associated diagnostic biomarkers.

Interpretation of these findings requires consideration of three limitations. First, although the identified bacterial proteins were recovered from the extracellular fraction, their routes of release remain unresolved. They may have entered the host-cell environment through conventional or noncanonical secretion, outer membrane vesicles, or bacterial lysis. We tried our best to prevent any lysis of bacteria and hence used cold water rather than Triton-X-100 and Saponin. We observed that Triton-X-100 killed *in vitro* a substantial number (up to 40%) of Ab. Saponin was expected to come in the way of convential proteomics. Sequential labelling identifies the period of protein synthesis independently of the mechanism of extracellular release. Accordingly, the term *released proteins* is more appropriate for these individual candidates until their export mechanisms are established. The observation that only 1.9% of protein groups identified under axenic conditions were recovered in the infection-associated extracellular fraction (Figure 4D; Supplementary Table S17) argues against simple equivalence between the two proteomes but does not exclude contributions from partial bacterial lysis. Second, although several proteins have known or predicted functions relevant to bacterial physiology and pathogenesis, their specific contributions to infection have not been experimentally established. Targeted gene deletion, complementation, and phenotypic characterization would be required to investigate these roles. Third, the labelling strain carried a pVRL2-derived expression plasmid and exhibited slower growth in minimal medium under strong induction than unmodified Ab7. A moderate growth burden associated with arabinose-induced expression from this vector has previously been reported (36). Although this may restrict labelling efficiency and proteomic coverage, the use of matched experimental conditions supports comparisons between the labelling groups. The enrichment of predicted signal peptides and outer-membrane proteins provides evidence of non-random composition, but neither these predictions nor the limited overlap with axenic fractions can quantify the contribution of bacterial lysis. Importantly, the cold-water recovery procedure was selected to minimize bacterial disruption; this methodological precaution strengthens sample interpretation without proving that all recovered bacterial proteins were actively secreted.

In addition to these biological limitations, two aspects of the proteomic analysis warrant consideration. First, including a broad range of BONCAT-associated modifications in the database search resulted in a small number of peptide-spectrum matches assigned modifications unrelated to the analogues used experimentally. Second, because gel fractions were obtained from infected A549 cultures, they contained both host- and bacterial-derived proteins. Searching these mixed samples against a bacterial-only database resulted in some host-derived peptides being incorrectly assigned to bacterial proteins. Both categories of potentially misleading identifications were excluded from the final analyses (Supplementary Tables S5 and S4). These observations emphasise the importance of restricting modification searches to experimentally relevant chemistries and using appropriate host–pathogen sequence databases when analysing infection-derived proteomic samples.

The methodological significance of our approach becomes clearer when considered alongside earlier applications of metabolic protein labelling during infection. Cell-selective analogue tagging has previously enabled the identification of bacterial proteins secreted during host infection (30,62), while AHA labelling has been employed to enrich newly synthesised bacterial proteins from infected host cells (63). Building on these approaches, our sequential AHA–ANL labelling strategy distinguishes bacterial proteins synthesised before host contact from those synthesised during infection. Importantly, analogue-containing peptides were identified directly without affinity enrichment, and the background frequency of analogue assignments was experimentally assessed. The nine proteins assigned to synthesis during co-culture therefore constitute a focused set of candidates for subsequent investigation through gene deletion, complementation, epitope tagging, and affinity-based recovery from infected cultures. Although their functions and routes of extracellular release remain to be established, their temporal identification provides information that cannot be obtained directly from transcript abundance or conventional steady-state protein measurements alone.

The translational value of the combined dataset, therefore, lies in a tiered candidate-selection strategy: proteins with ANL-modified peptides offer evidence of synthesis during host contact, whereas proteins recovered only through conventional identification broaden the pool of potential diagnostic antigens. Candidates could subsequently be prioritised using their reproducibility across biological replicates, predicted extracellular accessibility, abundance, and detectability in clinically accessible specimens. Establishing their diagnostic utility will require independent validation in patient samples, particularly because detection in the A549 infection model does not establish circulation or stability in neonatal blood.

## Acknowledgments

We sincerely thank M. Lucidi and colleagues for kindly providing the pVRL2 plasmid used to construct the recombinant vector for MetRSᴺᴸᴸ expression in Ab7. We acknowledge the Translational Health Science and Technology Institute (THSTI), Faridabad, India, for providing institutional support and research infrastructure for this study. We thank Surjeet Yadav for all his laboratory maintenance-related efforts and the administrative, purchase, and finance teams of THSTI for their support. We thank the Council of Scientific and Industrial Research for the fellowship to M.I. We also acknowledge the Institutional Biosafety Committee approval (IBSC # 288/2021) for procuring the required Ab isolates from the local hospital.

## Use of AI

Python scripts were used to visualise the data in Fig. 4. These were drafted with assistance from Claude (Anthropic, 2026) and subsequently reviewed, modified, and executed by the authors. Grammarly and ChatGPT (OpenAI) were used for English-language editing and to improve the clarity of the final draft of the manuscript.

## Conflicts of Interest

The authors declare no conflicts of interest, including financial interests, related to this work.

---

**Supplementary Table S1:**
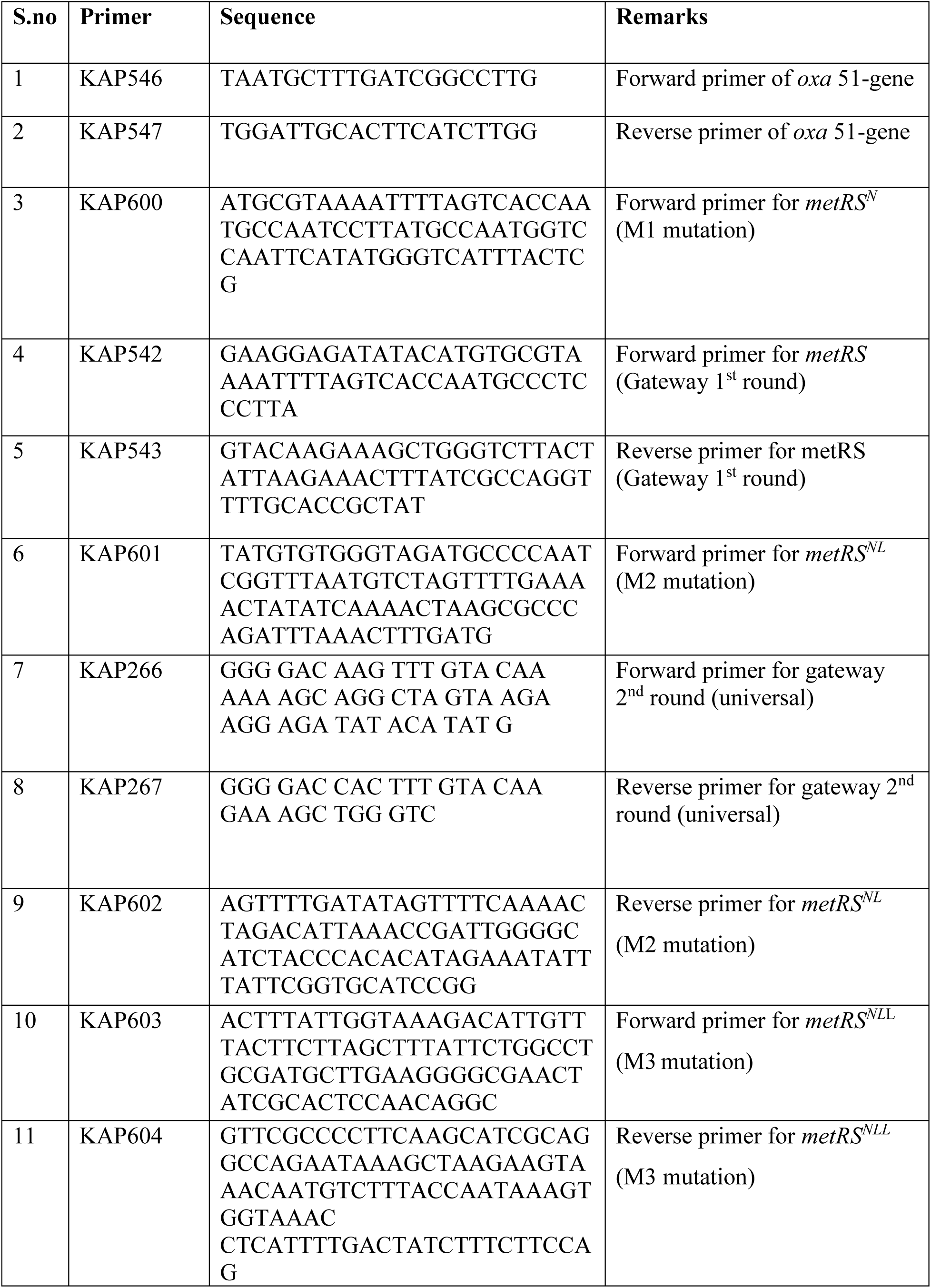
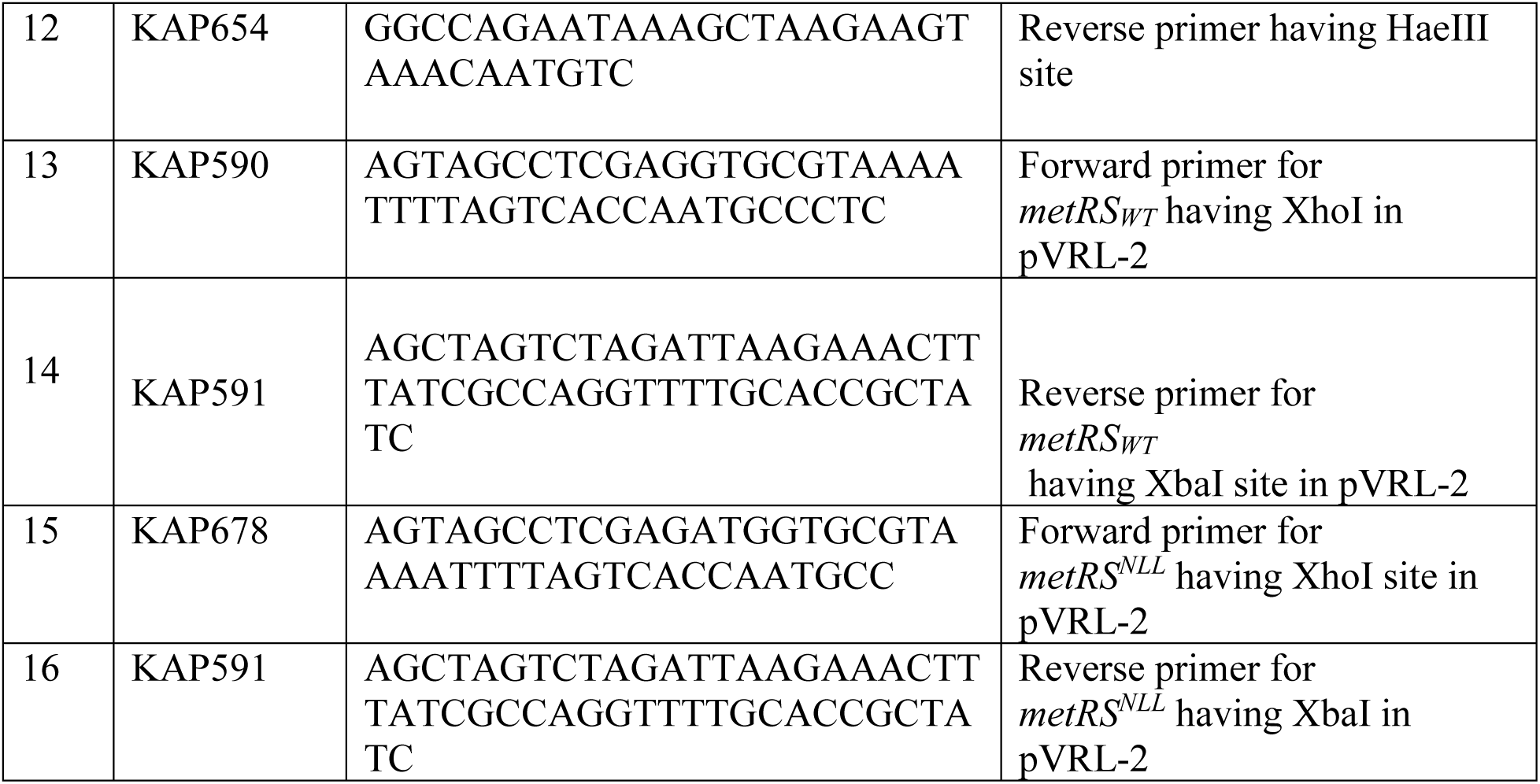
List of primers used in this study.

**Supplementary Table S2:** List of plasmids used in this study.

| S.no | Construct | Plasmid name | Reference |
| --- | --- | --- | --- |
| 1 | Shuttle vector (*E. coli* & Ab) | pVRL-2 | Lucidi M et al. 2018 |
| 2 | Gateway entry vector | pDONR-221 | Thermo Fisher<br>Scientific |
| 3 | Standard cloning vector | pBluescript KS (+) | Addgene |
| 4 | *metRS<sup>N</sup>* + pDONR-221 | pMI1 | This study |
| 5 | *metRS<sup>NL</sup>* + pDONR-221 | pMI2 | This study |
| 6 | *metRS<sup>NLL</sup>* in pBluescript KS (+) | pMI3 | This study |
| 7 | *metRS<sup>NLL</sup>* in pVRL-2 | pMI4 | This study |
| 8 | *metRS<sup>Wt</sup>* in pVRL-2 | pMI5 | This study |
| 9 | *metRS<sup>Wt</sup>* + pET28a | pMI6 | This study |

**Supplementary Figure 1:**
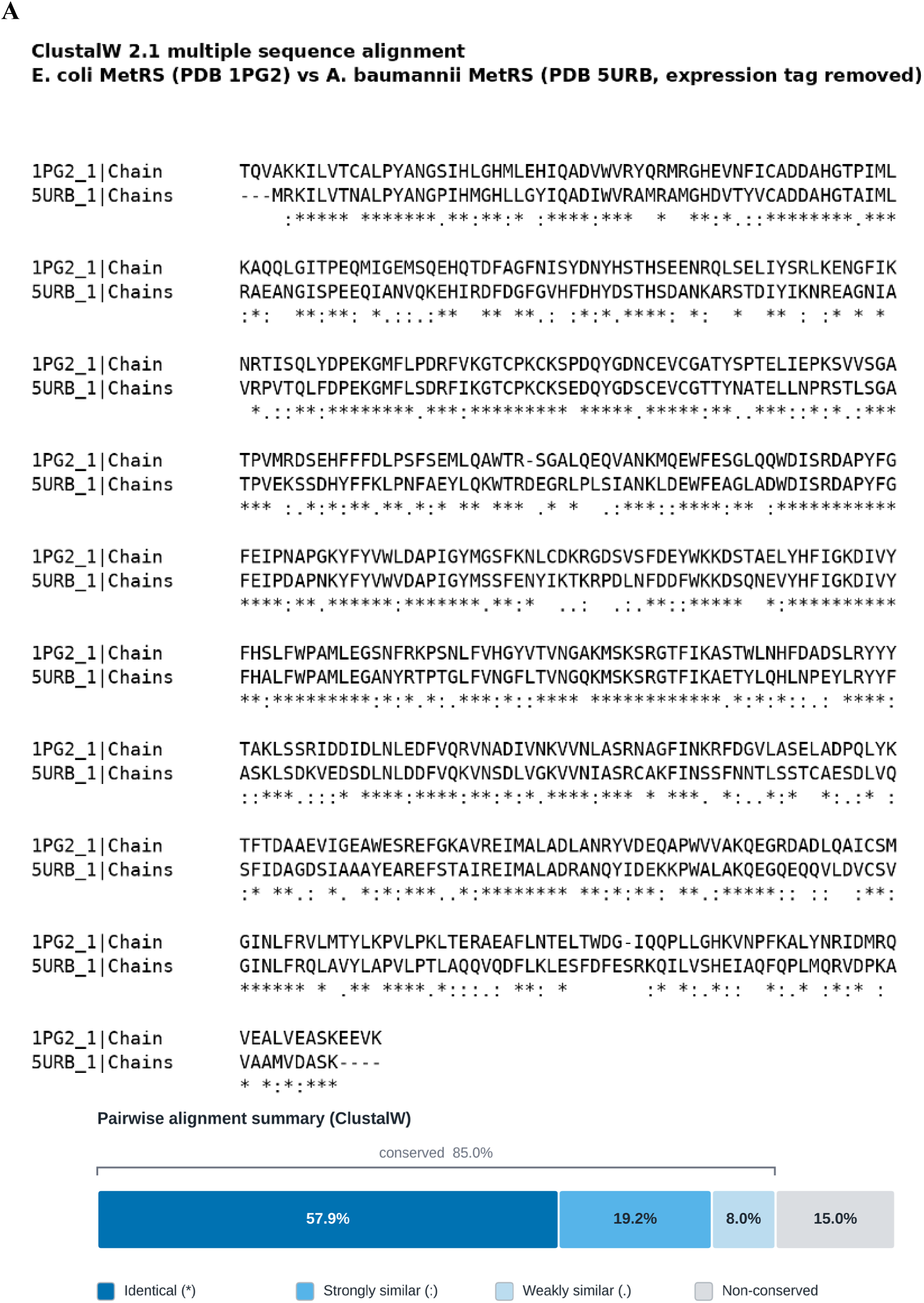
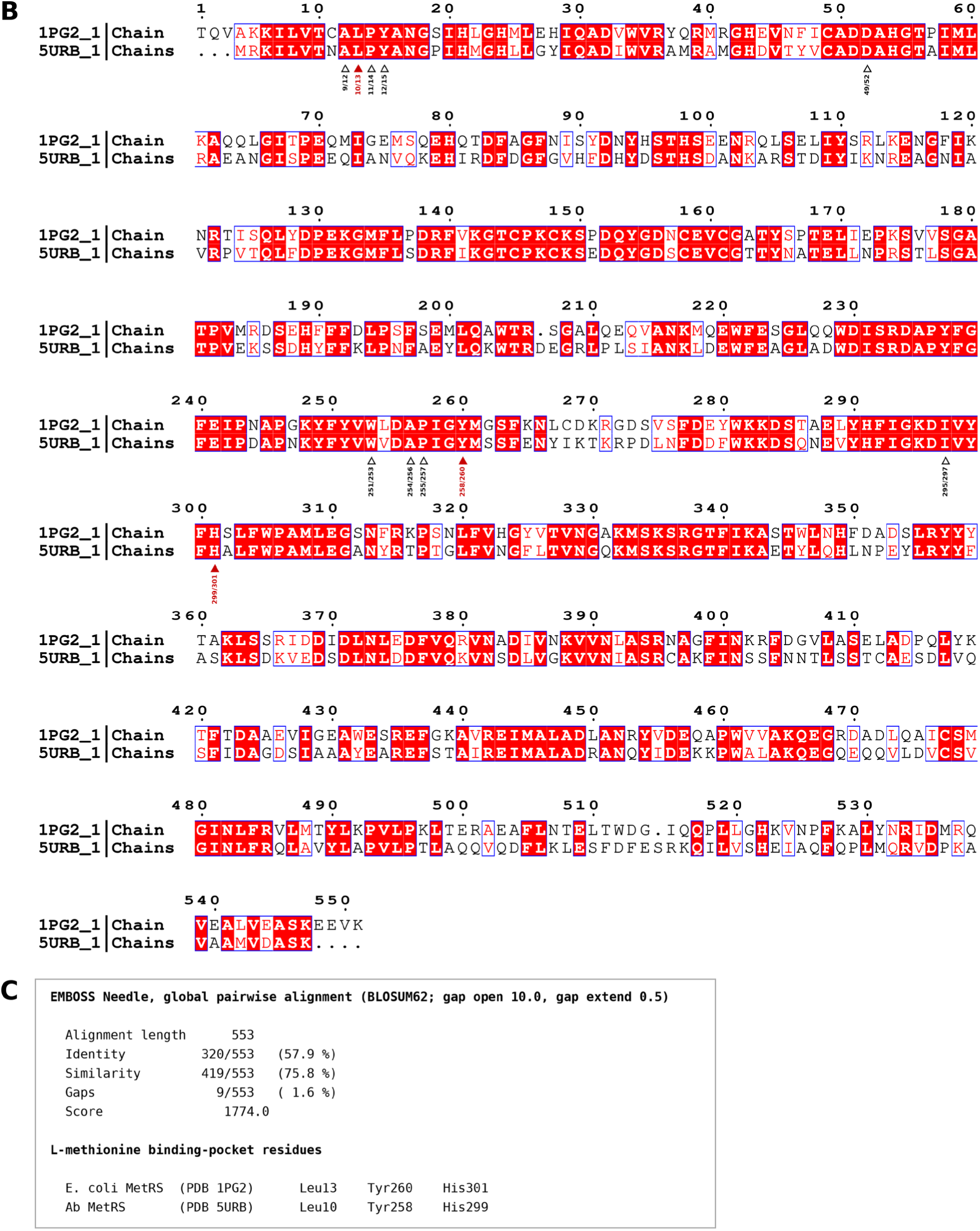
Sequence alignment of Ab and *E. coli* methionyl-tRNA synthetases (MetRS) reveals conservation of L-methionine-binding residues. **(A):** Summary of the ClustalW pairwise sequence alignment of Ab and *E. coli* MetRS proteins, showing 57.9% identical (*), 19.2% strongly similar (:), 8.0% weakly similar (.), and 15.0% non-conserved residues, corresponding to an overall sequence conservation of 85.0%. **(B):** Full-length sequence alignment of Ab MetRS (PDB: 5URB; N-terminal expression tag removed) and *E. coli* MetRS (PDB: 1PG2), generated using ClustalW and visualized with ESPript 3. Identical residues are represented by **white** letters on a **red** background, whereas similar residues are shown as **red** letters within **blue-outlined** boxes. **Triangles** beneath the alignment indicate **11** residues located within **4.5 Å** of the bound **L-methionine**, with residue positions annotated for Ab and *E. coli*. Filled red triangles identify the three residues selected for substitution in the engineered MetRSᴺᴸᴸ variant. These correspond to **Leu10, Tyr258,** and **His299** in Ab MetRS and **Leu13, Tyr260,** and **His301** in *E. coli* MetRS. An independent global pairwise alignment using EMBOSS Needle (BLOSUM62; gap-opening penalty, 10.0; gap-extension penalty, 0.5) identified 320 identical residues across 553 aligned positions (57.9% identity), with 1.6% gaps, consistent with the ClustalW alignment. The sequence ruler follows the residue numbering of *E. coli* MetRS (PDB: 1PG2).

**Supplementary Figure 2:**
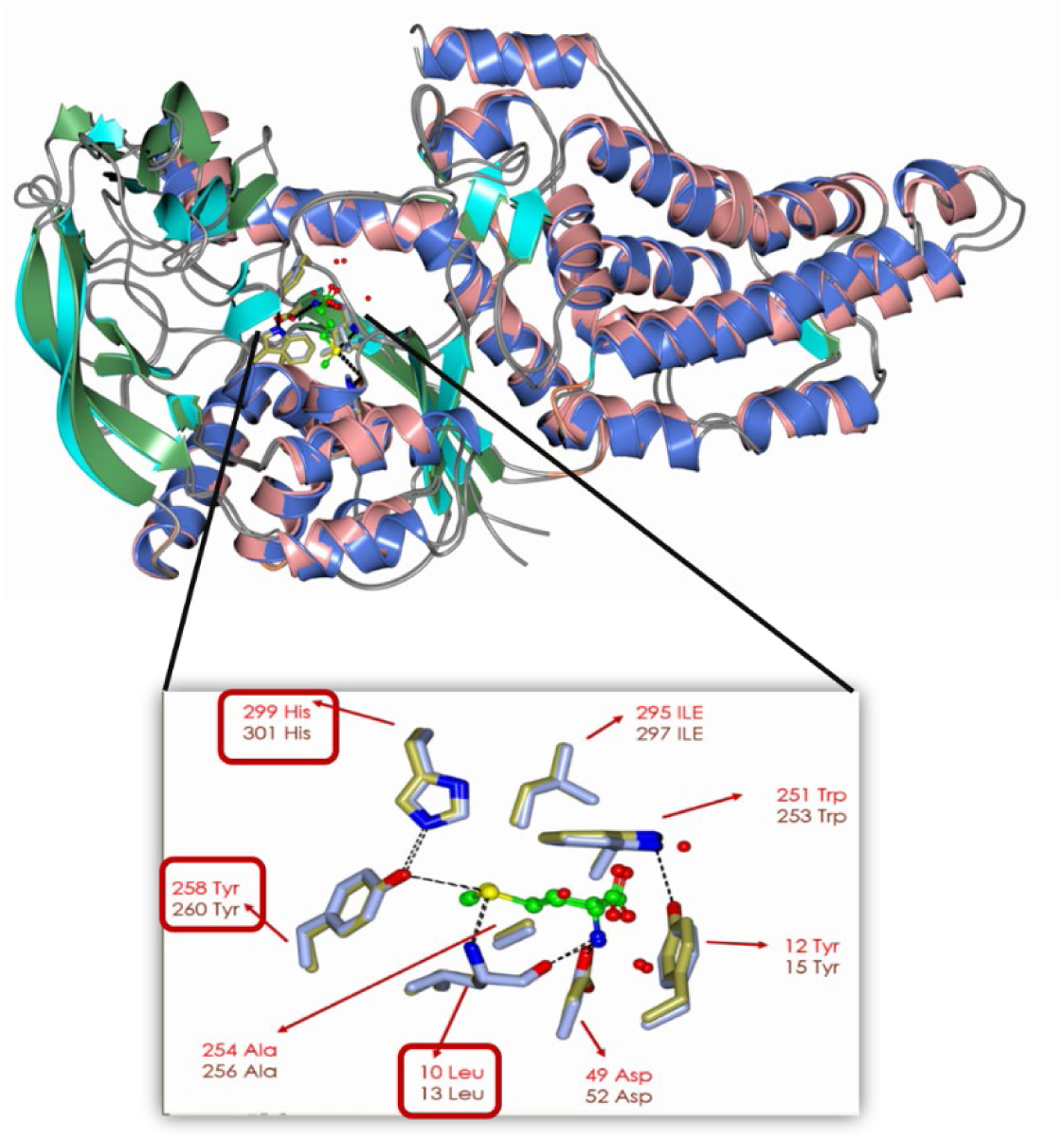
Structural conservation of the L-methionine-binding pocket in Ab and *E. coli* methionyl-tRNA synthetases (MetRS). Structural superimposition of Ab MetRS (PDB: 5URB; X-ray resolution, 1.90 Å; L-methionine-bound) and *E. coli* MetRS (PDB: 1PG2; X-ray resolution, 1.75 Å; methionine- and adenosine-bound), performed using WinCoot. **Upper panel:** Superimposition of the two MetRS structures, illustrating their overall structural similarity **Lower panel:** Enlarged view of the conserved L-methionine-binding pocket with bound L-methionine represented in ball-and-stick format. Ab residues are in grey, *E. coli* residues are in gold, and hydrogen bonds are shown as dashed lines. Eight of the 11 residues lining the binding pocket are indicated (all 11 are identical), with residue positions annotated as Ab/*E. coli*: Leu10/Leu13, Tyr12/Tyr15, Asp49/Asp52, Trp251/Trp253, Ala254/Ala256, Tyr258/Tyr260, Ile295/Ile297, and His299/His301. Boxed labels highlight the three residues substituted in the engineered Ab MetRSᴺᴸᴸ variant: L10N, Y258L, and H299L.

**Supplementary Figure 3:**
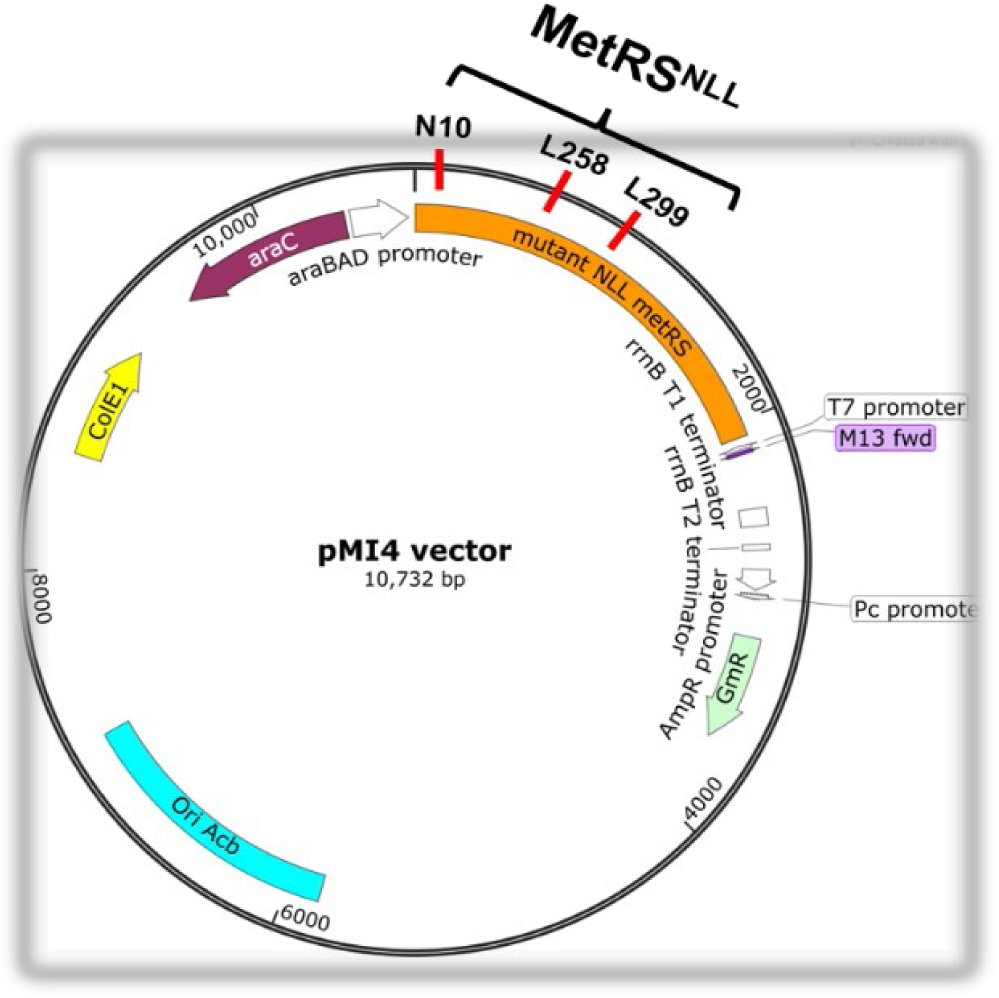
Schematic representation of the pMI4 shuttle plasmid encoding the engineered *A. baumannii* MetRSᴺᴸᴸ variant for ANL incorporation. Circular plasmid map of the pMI4 shuttle vector carrying the engineered Ab methionyl-tRNA synthetase (MetRSᴺᴸᴸ) gene, containing three amino acid substitutions within the L-methionine-binding pocket: L10N (M1), Y258L (M2), and H299L (M3). Expression of the MetRSᴺᴸᴸ variant is under the arabinose-inducible pBAD promoter, with induction achieved by using 2% arabinose. The plasmid map was generated using SnapGene Viewer.

**Supplementary Figure 4:**
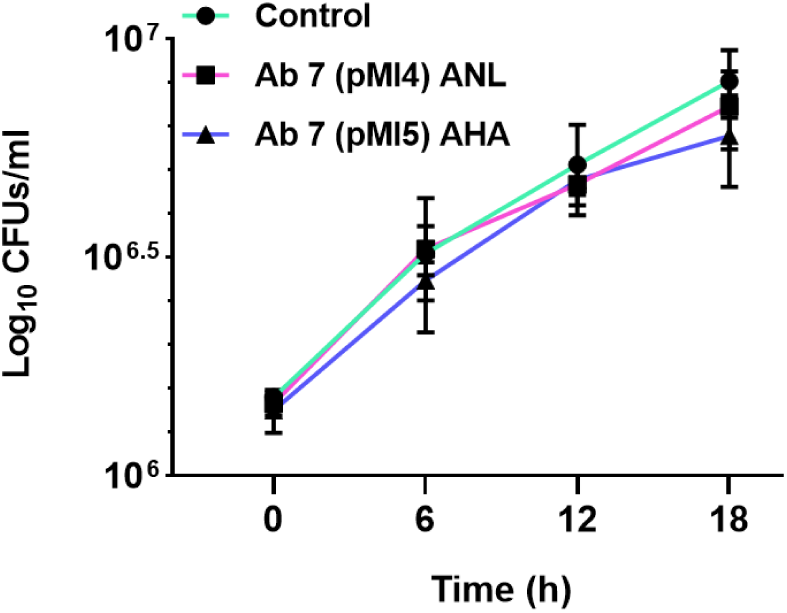
Incorporation of ANL and AHA does not affect the growth kinetics of Ab7. Growth kinetics of Ab7 expressing wild-type MetRS (MetRS^LYH^) or engineered MetRSᴺᴸᴸ in the presence of AHA and ANL, respectively. Primary cultures were grown in LB medium, washed twice with M9 minimal medium, and subcultured in M9 medium at an initial OD₆₀₀ of 0.005. Upon reaching an OD₆₀₀ of 0.1, MetRSᴺᴸᴸ expression was induced with 2% arabinose, supplemented with methionine biosynthesis inhibitory amino acids (MBIAA) and 1 mM ANL. Cultures expressing MetRS^LYH^ were supplemented with 0.25 mM AHA. Corresponding cultures grown without ANL or AHA served as controls. Bacterial growth was monitored by enumerating CFUs at different time points during the logarithmic growth phase. CFU counts (Y-axis) were plotted against sampling time (X-axis). Comparable growth kinetics across labelled and unlabelled cultures indicate that incorporation of ANL or AHA did not appreciably affect Ab7 growth under the experimental conditions.

**Supplementary Figure 5:**
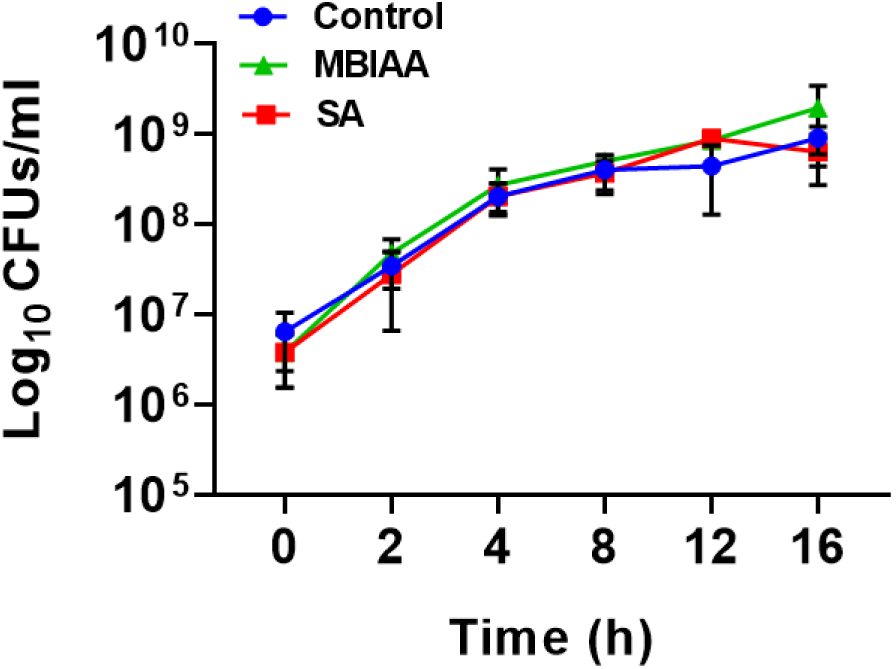
Sodium acetate and methionine biosynthesis inhibitory amino acids (MBIAA) do not affect the growth kinetics of WT Ab7. Growth kinetics of WT Ab7 cultured in the presence of 8 mM Sodium acetate or Methionine biosynthesis inhibitory amino acids (MBIAA). Ab7 was initially grown in LB medium, washed three times with M9 minimal medium, and subsequently subcultured into M9 medium supplemented with sodium acetate or MBIAA at an initial OD₆₀₀ of 0.005. Bacterial growth was monitored by enumerating colony-forming units (CFUs) at different time points. CFU counts were plotted on a logarithmic scale (Y-axis) against sampling time (X-axis). Growth kinetics were evaluated using three independent biological replicates. Neither treatment appreciably affected Ab7 growth under the experimental conditions.

**Supplementary Figure 6:**
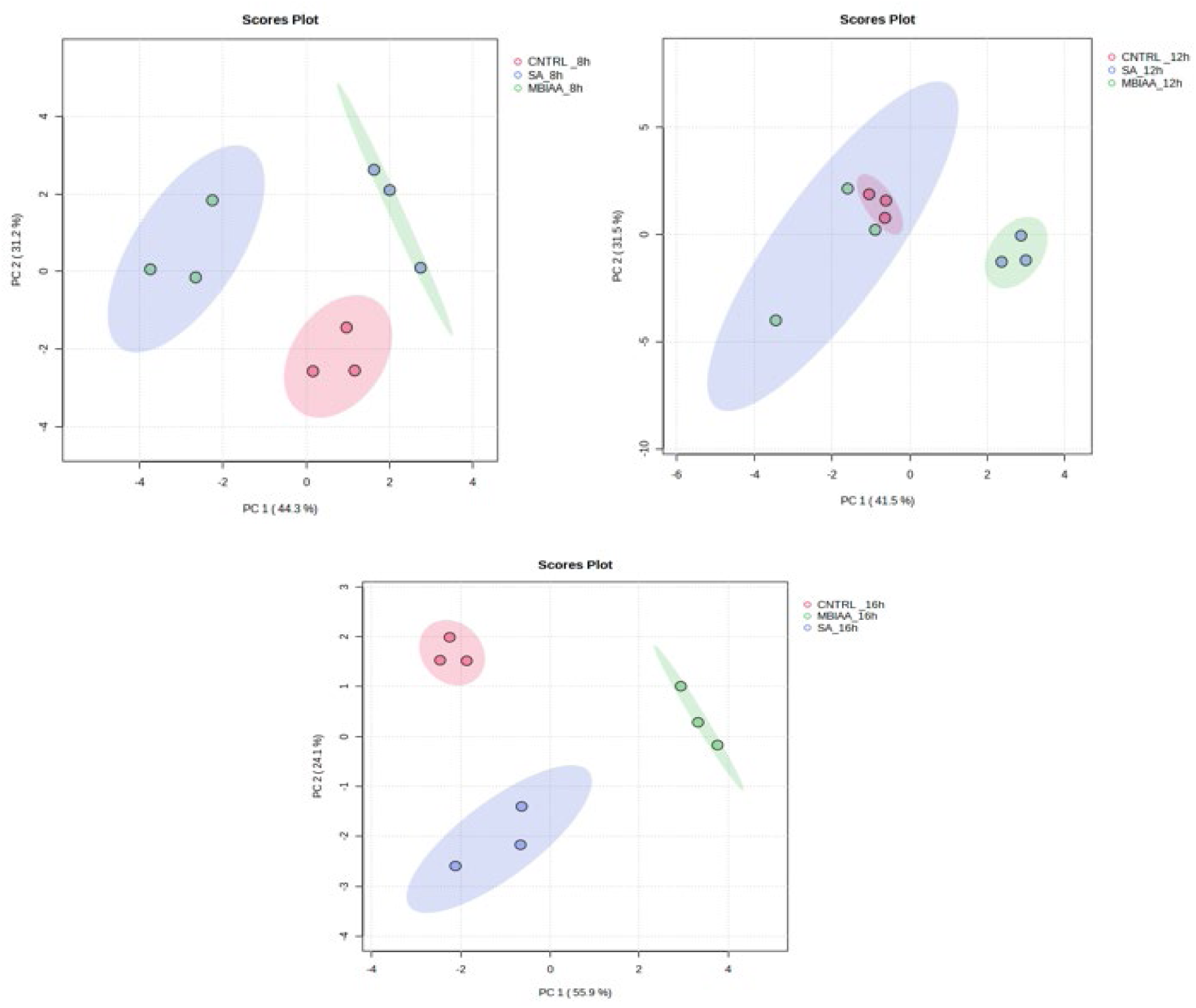
Principal component analysis (PCA) of the metabolomic profiles of Ab7 following Sodium acetate and MBIAA treatment. Principal component analysis (PCA) score plots of metabolomic profiles obtained from 02 untreated control, sodium acetate-treated, and methionine biosynthesis inhibitory amino acid 03 (MBIAA)-treated Ab7 cultures at 8, 12, and 16 h. PCA was performed separately at each time 04 point to assess differences in the overall metabolic profiles among the three experimental 05 groups.

**Supplementary Figure 7:**
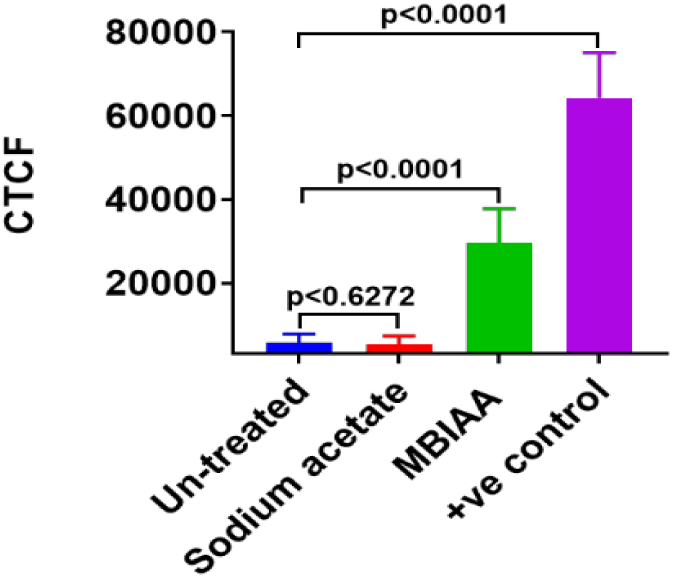
Quantitative fluorescence analysis confirms efficient ANL incorporation by Ab MetRSᴺᴸᴸ in the presence of MBIAA. Corrected total cell fluorescence (CTCF) was quantified to compare ANL incorporation mediated by engineered MetRSᴺᴸᴸ in the presence of MBIAA. Statistical significance was determined using an unpaired Student’s *t*-test. ns, not significant (*P* = 0.6272); ****, *P* < 0.0001.

**Supplementary Figure 8:**
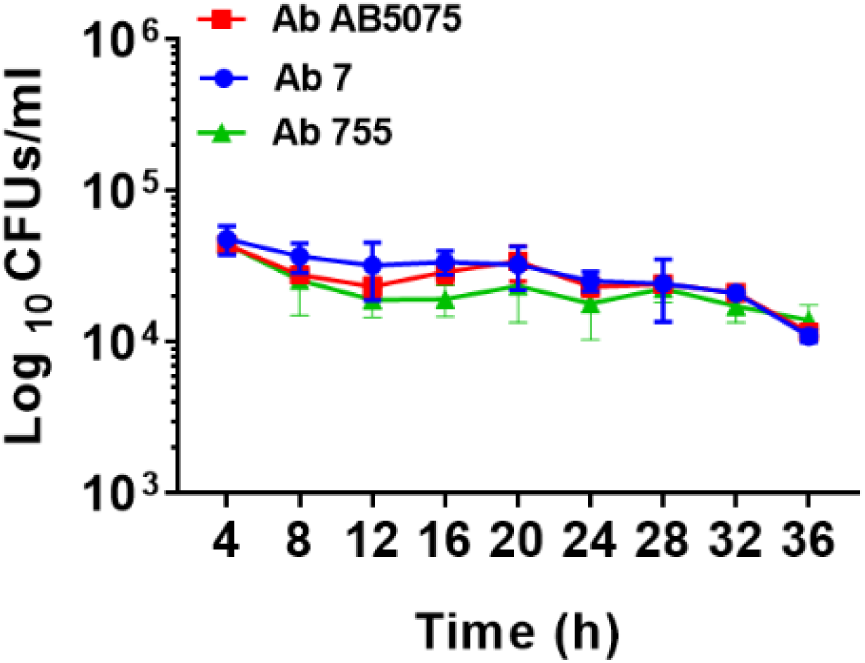
Ab7 exhibits intracellular survival kinetics comparable to those of the hypervirulent strain AB5075 and clinical isolate Ab755 in A549 lung epithelial cells. Intracellular survival of Ab7 and Ab755 (clinical isolates from our laboratory collection) was compared with that of the hypervirulent strain AB5075, originally isolated from a patient with tibial osteomyelitis. Human lung epithelial cells (A549) were infected with each strain at a multiplicity of infection (MOI) of 1000 for 3 h, followed by treatment with colistin (8 µg/mL) for 1 h to eliminate extracellular bacteria. Infected A549 cells were subsequently lysed using cold water, and intracellular bacterial survival was assessed by enumerating CFUs (Y-axis) at different post-infection time points (X-axis). Ab7 remained recoverable from A549 cells for up to 24 h post-infection, exhibiting survival kinetics comparable to those of AB5075 and Ab755.

**Supplementary Figure 9.**
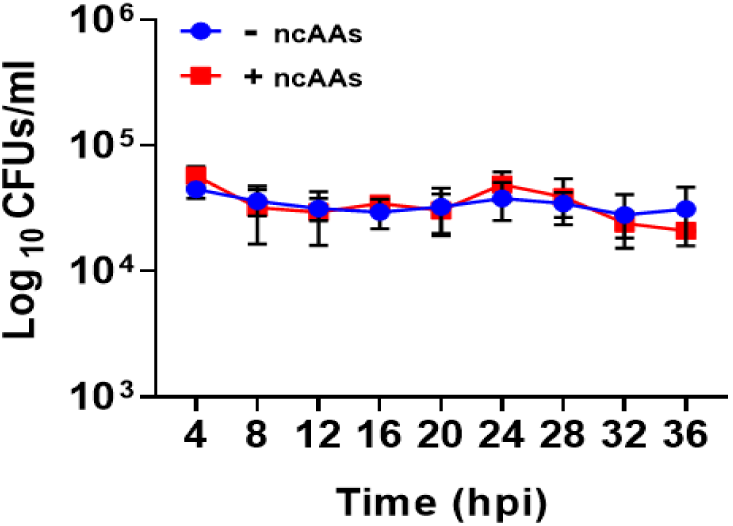
Incorporation of non-canonical amino acids (ncAAs) does not affect the intracellular survival of Ab7 in A549 epithelial cells. Intracellular survival of Ab7 expressing wild-type MetRS (MetRS^LYH^) or engineered MetRSᴺᴸᴸ was evaluated following infection of human lung epithelial cells (A549) at a multiplicity of infection (MOI) of 1: 1000. Bacterial cultures were grown in M9 minimal medium with (+ncAAs) or without (−ncAAs) AHA, followed by infection of A549 cells in the presence or absence of ANL, as indicated. At 3 h post-infection (hpi), infected cells were washed three times with PBS and treated with antibiotics to eliminate extracellular bacteria. At the indicated time points, A549 cell monolayers were washed and lysed, and intracellular bacterial counts determined by plating serial dilutions on LB agar. CFUs (Y-axis) were plotted against time post-infection (X-axis). Comparable intracellular bacterial counts in labelled and unlabelled conditions indicate that ncAA incorporation did not appreciably affect Ab7 survival within A549 cells.

**Supplementary Figure 10.**
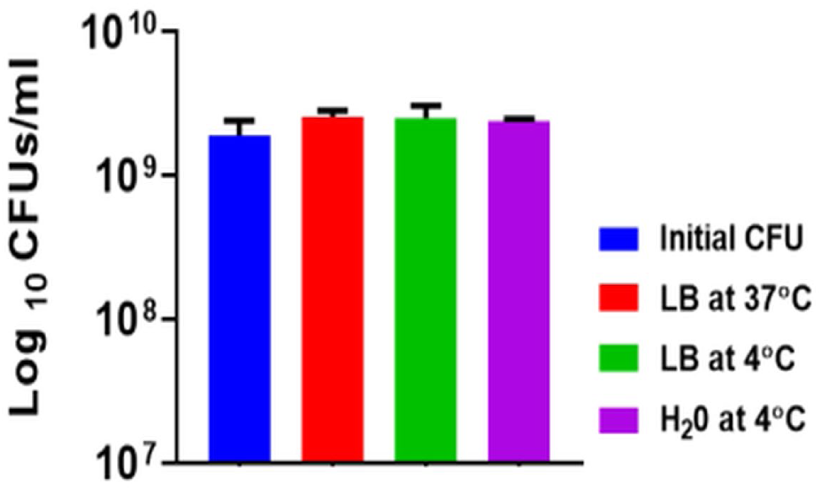
Hypotonic treatment with cold water does not affect the viability of Ab7. 1 mL of late logarithmic-phase Ab7 cultures (OD₆₀₀ ≈ 3.0) were harvested by centrifugation and resuspended in 1 mL of LB medium at 37 °C, LB medium at 4 °C, or sterile water at 4 °C for 1 h. An untreated bacterial suspension served as the input control (initial CFU). Following treatment, viable bacterial counts were determined by serial dilution and plating on LB agar. Comparable CFU counts across all conditions demonstrated that exposure to cold sterile water did not appreciably affect bacterial viability, supporting its use for hypotonic lysis of infected A549 cell monolayers without compromising the recovery of viable bacteria. Data are presented as mean ± standard deviation (SD) from three independent biological replicates.

**Supplementary Figure 11:**
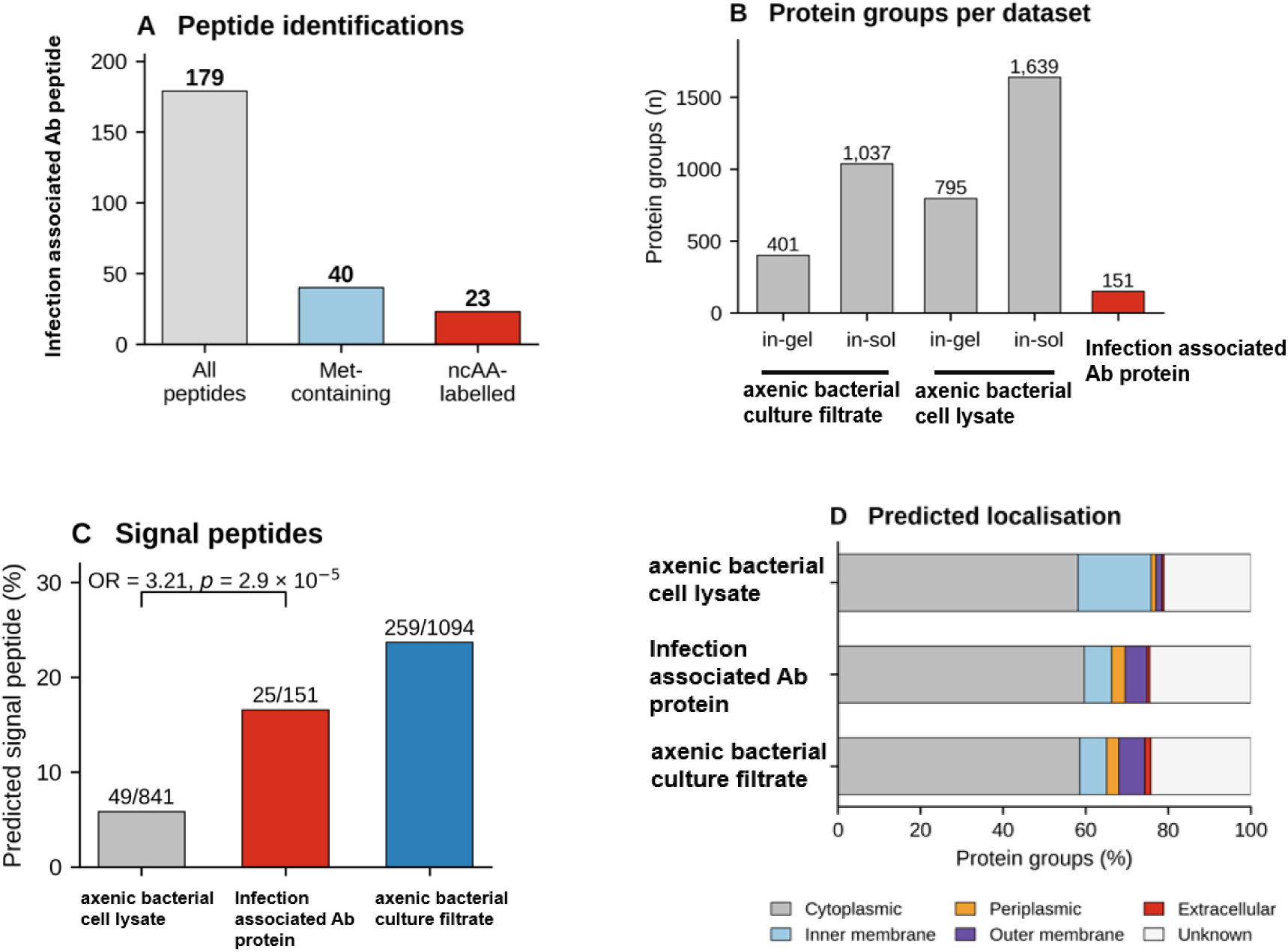
Summary of protein identification, signal peptide prediction, and subcellular localization. **(A)** Number of peptides retained from the infection-associated Ab protein after exclusions (n = 179), including methionine-containing peptides (n = 40), which permit detection of analogue incorporation, and peptides with site-localized amino acid analogues (n 83 = 23). Corresponding control data are provided in Supplementary Table S6. (B) Number of protein groups identified in each dataset. Axenic samples were analyzed using both in-gel and in-solution digestion, whereas infection-associated Ab protein samples were analyzed using in-gel digestion only. (C) Proportion of identified protein groups containing predicted signal peptides, as determined using SignalP 6.0. Numbers above the bars indicate the number of signal peptide-positive protein groups relative to the total number analyzed. Statistical significance was assessed using a two-sided Fisher’s exact test. (D) Predicted subcellular localization of identified protein groups using PSORTb v3.0.2. “Axenic bacterial cell lysate” denotes protein groups detected in the axenic bacterial cell lysate but not in the axenic bacterial culture filtrate.

**Supplementary Figure 12:**
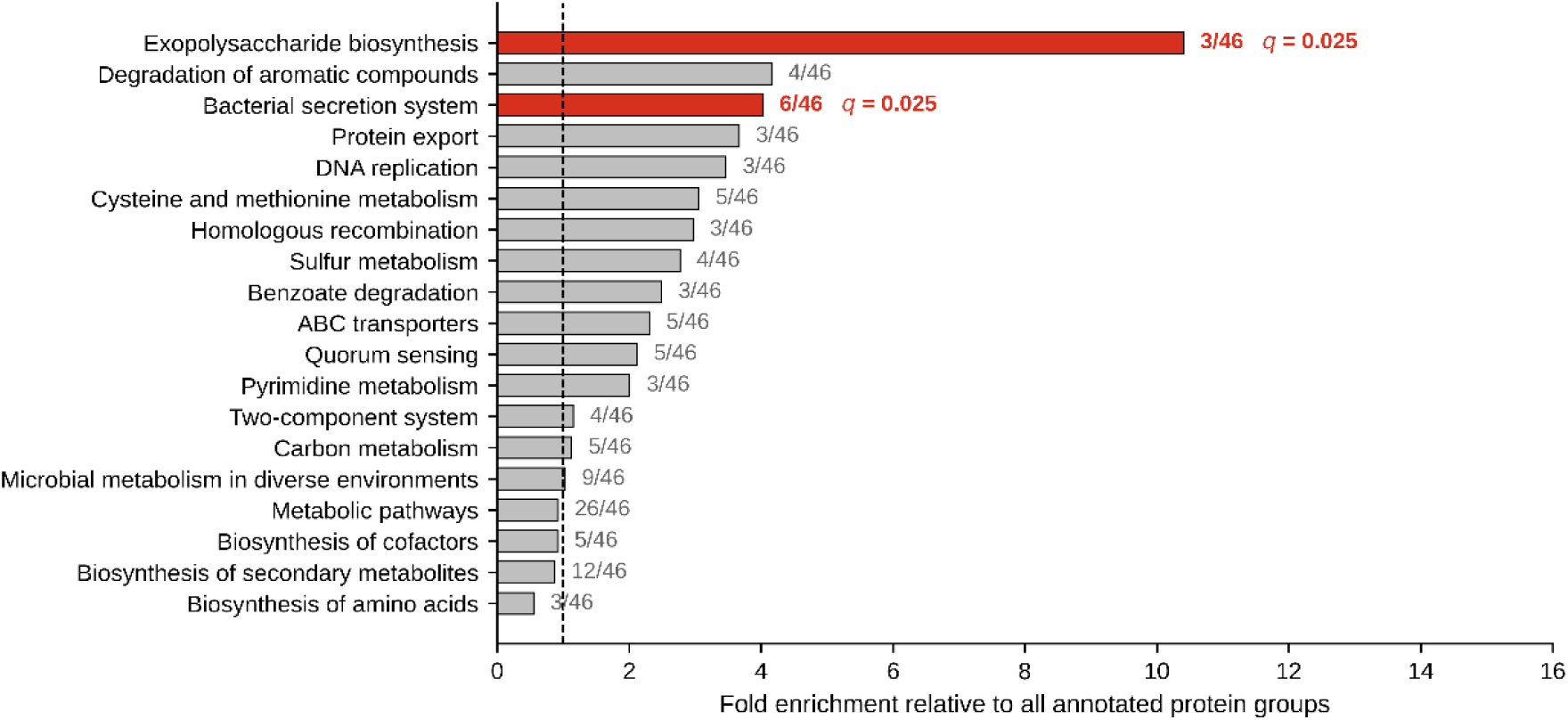
KEGG pathway enrichment analysis of infection-associated Ab protein groups. All 19 pathways that met the testing threshold are shown, ranked by fold enrichment relative to the background of all annotated protein groups. Numbers indicate the proportion of protein groups associated with each pathway relative to the total number of annotated protein groups in the 00 test set. Red bars indicate pathways significant after Benjamini–Hochberg correction (q < 01 0.05), whereas grey bars indicate non-significant pathways. The dashed line represents no 02 enrichment (fold enrichment = 1).

## Notes

### Competing Interest Statement

The authors have declared no competing interest.

### Summary of Updates

1. To specify that Dr. Mohd Ilyas is the first author and co-corresponding author. The # symbol has been inserted to represent the same. 2. The star has been indicated for the Main Corresponding Author. 3. Ruchi Tomar is changed to Ruchi Tanwer. The last name is now correct 4. Saifullah Mohd is changed to Saifullah Muzaffar Husain. The middle name and last name is now correct. Typically, many Indians do not have a last name until they marry. But given that bioRxiv requires authors to specify their last names, Ms. Ruchi and Mr. Saifullah were asked to provide their last names. I, the corresponding author, misspelled Ms. Ruchi's last name; it has now been corrected in the revised manuscript. Mr. Saifullah requested that his last name be changed from Mohd to Muzaffar Husain (his father's name), and the change was approved. Nothing else is altered in the manuscript, the supplementary doc, and Tables 3-19.

